# A tissue-engineered human trabecular meshwork hydrogel for advanced glaucoma disease modeling

**DOI:** 10.1101/2020.07.31.229229

**Authors:** Haiyan Li, Tyler Bagué, Alexander Kirschner, Robert W. Weisenthal, Alison E. Patteson, Nasim Annabi, W. Daniel Stamer, Preethi S. Ganapathy, Samuel Herberg

## Abstract

**Purpose:** Abnormal human trabecular meshwork (HTM) cell function and extracellular matrix (ECM) remodeling contribute to HTM stiffening in primary open-angle glaucoma (POAG). Most current cellular HTM model systems do not sufficiently replicate the complex native three dimensional (3D) cell-ECM interface, which makes them less than ideal to investigate POAG pathology. Tissue-engineered protein-based hydrogels are ideally positioned to overcome shortcomings of current models. Here, we report a novel biomimetic HTM hydrogel and test its utility as a POAG disease model.

**Methods:** HTM hydrogels were engineered by mixing normal donor-derived HTM cells with collagen type I, elastin-like polypeptide and hyaluronic acid, each containing photoactive functional groups, followed by UV light-activated free-radical crosslinking. Glaucomatous conditions were induced with dexamethasone (DEX), and therapeutic effects of the Rho-associated kinase (ROCK) inhibitor Y27632 on cytoskeletal organization and tissue-level function, contingent on HTM cell-ECM interactions, were assessed.

**Results:** DEX exposure increased HTM hydrogel contractility, f-actin and alpha smooth muscle actin abundance and rearrangement, ECM remodeling, and fibronectin and collagen type IV deposition, all contributing to HTM hydrogel condensation and stiffening consistent with recent data from normal vs. glaucomatous HTM tissue. Y27632 treatment produced precisely the opposite effects and attenuated the DEX-induced pathologic changes, resulting in HTM hydrogel relaxation and softening.

**Conclusions:** We have developed a biomimetic HTM hydrogel model for detailed investigation of 3D cell-ECM interactions under normal and simulated glaucomatous conditions. Its bidirectional responsiveness to pharmaceutical challenge and rescue suggests promising potential to serve as screening platform for new POAG treatments with focus on HTM biomechanics.

## Introduction

The human trabecular meshwork (HTM), located in the iridocorneal angle, is an avascular connective tissue with filtering features and complex architecture ^1^. Its main function is to control the resistance to drainage of aqueous humor (AH) via the conventional outflow pathway to maintain normal intraocular pressure (IOP) ^2^. Most of the AH outflow resistance is localized to the deepest juxtacanalicular tissue (JCT) region ^3^, where a few discontinuous layers of HTM cells reside in and interact with a dense amorphous extracellular matrix (ECM). A variety of ECM structural and organizational components have been identified within the JCT; e.g., non-fibrillar and fibrillar collagens, elastic fibrils, and basement membrane-like materials. The narrow space between HTM cells and ECM fibers is filled with a ground substance rich in proteoglycans and the glycosaminoglycan hyaluronic acid (HA) ^3–7^.

HTM cells play an essential role in modulating AH outflow resistance by controlling the production of contraction forces and the secretion/degradation of ECM proteins to support tissue homeostasis ^8^. As such, the reciprocity between HTM cells and their ECM is critical for normal tissue function and IOP maintenance within narrow margins in the healthy eye. Dysregulation of HTM cells can cause elevated ECM deposition as well as HTM cell and tissue contraction, which jointly contribute to HTM stiffening, ultimately leading to increased AH outflow resistance and elevated IOP ^9,10^. Prolonged elevation of IOP causes pathologic distension and compression of the HTM and further tissue stiffening. Consequently, the stiffened HTM negatively affects IOP and HTM cell function in a feed-forward loop ^11–13^. Since elevated IOP is a primary risk factor for glaucoma, this pathologic process leads to retinal ganglion cell damage resulting in the irreversible loss of vision that is characterized as primary open-angle glaucoma (POAG), the most common form of glaucoma ^14–18^.

Elevated IOP is the only modifiable risk factor for POAG, and management of the disease is focused on IOP lowering ^16^. First line POAG medications do not specifically target the diseased HTM; instead, they lower IOP by either increasing AH outflow bypassing the HTM altogether, or by decreasing AH production ^19,20^. In the United States, only the recently approved Rho-associated kinase (ROCK) inhibitor netarsudil ^21^ directly aims at the stiffened HTM to increase AH outflow via cell/tissue relaxation ^22–26^. However, various adverse events at the ocular surface in some patients have been reported during the drug’s brief window of availability ^27–29^, raising concerns over its routine use. This highlights the clear unmet need for new/improved POAG treatment strategies. Targeting ECM mechanics by reversing tissue stiffening to limit pathological progression is an emerging therapeutic approach in other diseases ^30^, with promising potential for POAG therapy.

A number of models are available to study HTM physiology and POAG pathophysiology; e.g., conventional two dimensional (2D) HTM cell monolayer cultures, perfusion-cultured anterior segments from postmortem human and animal eyes, or animal models of different (large and small) species ^31–35^. While each of these model systems has proven strengths, they cannot reliably and efficiently determine the relative contributions of HTM cells and their ECM to the onset and progression of glaucomatous HTM stiffening. This has hampered advances in the mechanistic study of this biomechanical element in POAG pathophysiology.

The interdisciplinary field of tissue engineering aims to produce functional biomimetic replicas of tissues of interest ^36^. HTM tissue engineering is still in its infancy; few studies have been reported that use bioengineered 3D HTM *in vitro* models. Constructs made of porous scaffolds (~20 µm thick) seeded with HTM cells were shown to partially mimic normal *in vivo* tissue function, including pharmacological induction/rescue of glaucomatous conditions ^37–39^. However, an argument could be made that this model is more akin to a 2D cell culture system (i.e., HTM cells grown on a flat culture substrate) rather than a true 3D model, as it cannot accurately mimic the 3D cell-ECM interface beyond ECM secreted by the HTM cells while grown atop the synthetic polymer scaffold. Another scaffold-based approach utilized porcine TM cells ^40^ or HTM cells ^41^ cultured atop a freeze-dried polymer matrix made of collagen and chondroitin sulfate and/or HA exhibiting aligned pores throughout the 3D construct (~3 mm thick). The scaffolds were shown to support TM cell proliferation into the porous matrix, and the bioengineered constructs displayed certain aspects of normal tissue function. Nevertheless, the study did not investigate the constructs’ behavior under induced glaucomatous conditions.

In addition to scaffold-based systems, viscoelastic hydrogels (i.e., water-swollen networks of polymers) are widely used in tissue engineering applications. They provide a simplistic version of the natural 3D tissue environment and allow for accurate *in vitro* modeling of cellular behaviors ^42–46^. To that end, Matrigel, the gelatinous protein mixture secreted by mouse sarcoma cells, has been used to fabricate a rudimentary 3D HTM hydrogel culture model ^47–49^. Significant drawbacks include Matrigel’s tumorigenic origin, diverse composition, and batch-to-batch variability affecting its biochemical and mechanical properties ^50^. Lastly, a recent study reported a shear-thinning peptide hydrogel with HTM cells seeded atop to engineer an injectable HTM implant with utility as *in vitro* model ^51^. However, this simple 20-residue peptide cannot faithfully recapitulate the complex ECM environment HTM cells interact with in the native tissue. Together, these studies highlight the need for more well-defined hydrogel systems to support relevant HTM modeling studies under normal and induced glaucomatous conditions.

Here, we report a novel tissue-engineered HTM hydrogel composed of normal donor-derived HTM cells and ECM biopolymers found in the native tissue, with focus on modeling the JCT region owing to both its critical role in regulating AH outflow resistance and IOP, and inextricable link to glaucomatous HTM stiffening. HTM hydrogels in different sizes/shapes were formed by mixing HTM cells with collagen type I, HA, and elastin-like polypeptide (ELP) - each containing photoactive functional groups - followed by photoinitiator-mediated short UV crosslinking. We then used proven pharmacological induction of glaucomatous conditions by treating HTM hydrogels with the corticosteroid dexamethasone (DEX) ^35^ and assessed the therapeutic effects of the ROCK inhibitor Y27632 ^52^, either as co- or sequential treatment, on cytoskeletal organization and tissue-level functional changes contingent on HTM cell-ECM interactions. We showed that DEX increased HTM hydrogel contractility, f-actin and alpha smooth muscle actin (αSMA) abundance and rearrangement, ECM remodeling, and fibronectin and collagen type IV deposition, all contributing to HTM hydrogel condensation and stiffening. Y27632 treatment had precisely opposite effects and was shown to prevent or rescue the DEX-induced pathologic changes, resulting in HTM hydrogel relaxation and softening. Together, we have developed a biomimetic HTM hydrogel model that allows for detailed investigation of 3D cell-ECM interactions under normal and simulated glaucomatous conditions. The bidirectional responsiveness of the HTM hydrogel model system to pharmaceutical challenge and rescue suggests promising potential to serve as a screening platform for new POAG treatments with focus on HTM cell/tissue biomechanics.

## Materials and Methods

### HTM cell isolation and culture

Human donor eye tissue use was approved by the SUNY Upstate Medical University Institutional Review Board (protocol #2018-0), and all experiments were performed in accordance with the tenets of the Declaration of Helsinki for the use of human tissue. Normal human trabecular meshwork (HTM) cells were isolated from healthy donor corneal rims discarded after transplant surgery, and cultured according to established protocols ^53,54^. Using an SMZ1270 stereomicroscope (Nikon Instruments, Melville, NY, USA), corneal rims were cut into wedges and the HTM tissue was grabbed by placing one tip of Dumont #5 fine-tipped forceps (Fine Science Tools, Foster City, CA, USA) into the Schlemm’s canal lumen and the other on top of the anterior HTM. The dissected strips of HTM tissue were digested for 20 min with 1 mg/ml collagenase (Worthington, Lakewood, NJ, USA) and 15 μl/ml human albumin (Sigma-Aldrich, St. Louis, MO, USA) in Dulbecco’s Phosphate Buffered Saline (DPBS; Gibco; Thermo Fisher Scientific, Waltham, MA, USA) at 37°C, placed into a single well of gelatin-coated (Sigma-Aldrich) 6-well culture plates (Corning; Thermo Fisher Scientific), and overlaid with glass coverslips to aid in tissue adherence. HTM tissue strips were cultured in low-glucose Dulbecco’s Modified Eagle’s Medium (DMEM; Gibco) containing 20% fetal bovine serum (FBS; Atlanta Biologicals, Flowery Branch, GA, USA) and 1% penicillin/streptomycin/glutamine (PSG; Gibco), and maintained at 37°C in a humidified atmosphere with 5% CO2. Fresh media was supplied every 2-3 days. Once confluent, HTM cells were lifted with 0.25% trypsin/0.5 mM EDTA (Gibco) and sub-cultured in DMEM with 10% FBS and 1% PSG. All studies were conducted using cells passage 3-7; the reference HTM cell strain HTM129 was isolated and characterized at Duke University by W.D.S. (Table 1).

**Table 1.** HTM cell strain information.

| ID | Sex | Age |
| --- | --- | --- |
| Reference (HTM129)* | Female | 75 |
| HTM01 | Female | 26 |
| HTM11 | Female | 47 |
| HTM12 | Female | 60 |
| *\*obtained from the Stamer laboratory at Duke University* |  |  |

### HTM cell characterization

HTM cells were seeded at 1×10^4^ cells/cm^2^ in 6-well culture plates or on glass coverslips in 24-well culture plates, and cultured in DMEM with 10% FBS and 1% PSG. HTM cell morphology and growth characteristics were monitored by phase contrast microscopy using an LMI-3000 Series Routine Inverted Microscope (Laxco; Thermo Fisher Scientific). Once confluent, HTM cells were treated with 100 nM dexamethasone (DEX; Fisher Scientific) or vehicle control (0.1% (v/v) ethanol) in DMEM with 1% FBS and 1% PSG for 4 d, and serum-and phenol red-free DMEM for 3 d. The HTM cell culture supernatants were collected and concentrated using Amicon® Ultra Centrifugal Filters (Millipore Sigma, Burlington, MA, USA) for immunoblot analysis. The monolayer HTM cells were processed for quantitative reverse transcription-polymerase chain reaction (qRT-PCR) and immunocytochemistry (ICC) analyses. DEX-induced myocilin (MYOC) upregulation in more than 50% of HTM cells was used as inclusion/exclusion criterion.

### Immunoblot analysis

Equal protein amounts (10 µg), determined by standard bicinchoninic acid assay (Pierce; Thermo Fisher Scientific), from concentrated HTM cell culture supernatants ± DEX at 7 d supplemented with Halt™ protease/phosphatase inhibitor cocktail (Thermo Fisher Scientific) in 6X loading buffer (Boston Bio Products, Ashland, MA, USA) with 5% beta-mercaptoethanol (BME; Fisher Scientific), were subjected to SDS-PAGE using NuPAGE™ 4-12% Bis-Tris Gels (Invitrogen; Thermo Fisher Scientific) at 200V for 50 min and transferred to 0.45 µm PVDF membranes (Millipore; Thermo Fisher Scientific). Membranes were blocked with 5% nonfat milk (Nestlé USA, Glendale, CA, USA) in tris-buffered saline with 0.2% Tween®20 (TBST; Thermo Fisher Scientific), and probed with a primary antibody against MYOC (anti-MYOC [MABN866] 1:2000; Sigma-Aldrich) followed by incubation with an HRP-conjugated secondary antibody (Cell Signaling, Danvers, MA, USA). Bound antibodies were visualized with the enhanced chemiluminescent detection system (Pierce) on autoradiography film (Thermo Fisher Scientific).

### Quantitative reverse transcription-polymerase chain reaction (qRT-PCR) analysis

Total RNA was extracted from HTM cells ± DEX at 7 d (N = 3 per group and donor) using PureLink RNA Mini Kit (Invitrogen). RNA concentration was determined with a NanoDrop spectrophotometer (Thermo Fisher Scientific). RNA was reverse transcribed using iScript™ cDNA Synthesis Kit (BioRad, Hercules, CA, USA). Fifty nanograms of cDNA were amplified in duplicates in each 40-cycle reaction using a CFX 384 Real Time PCR System (BioRad) with annealing temperature set at 60°C, Power SYBR™ Green PCR Master Mix (Thermo Fisher Scientific), and custom-designed qRT-PCR primers for MYOC and Glyceraldehyde 3-phosphate dehydrogenase (GAPDH) (IDT, Coralville, IA, USA; Table 2). Transcript levels were normalized to GAPDH, and DEX-induced MYOC mRNA fold-changes calculated relative to vehicle controls using the comparative CT method ^55^.

**Table 2.**
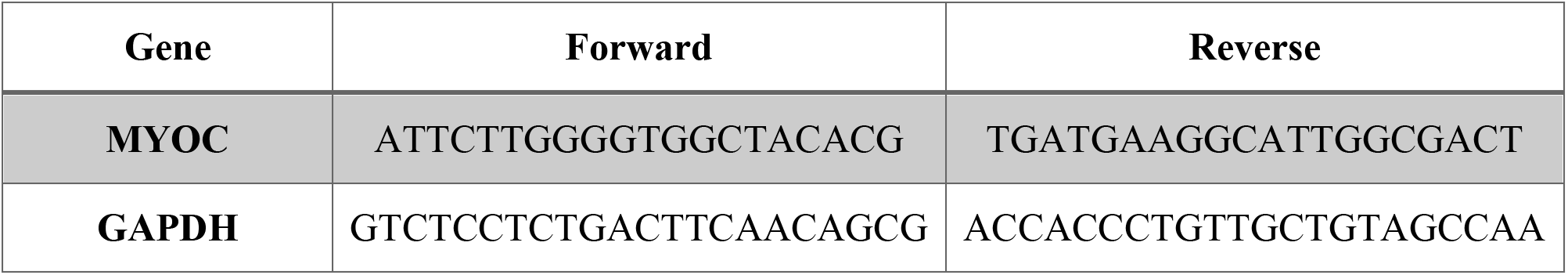
qRT-PCR primer information.

### Immunocytochemistry analysis

HTM cells ± DEX at 7 d were fixed with 4% paraformaldehyde (PFA; Thermo Fisher Scientific) at room temperature for 10 min, permeabilized with 0.5% Triton™ X-100 (Thermo Fisher Scientific), blocked with blocking buffer (BioGeneX, Fremont, CA, USA), and incubated with primary antibodies against MYOC (anti-MYOC [ab41552] 1:200; Abcam, Cambridge, MA, USA) or alpha B-Crystallin (anti-CRYAB [ab13497] 1:200; Abcam) followed by incubation with Alexa Fluor® 488-conjugated secondary antibodies (Abcam). Nuclei were counterstained with 4’,6’-diamidino-2-phenylindole (DAPI; Abcam). Coverslips were mounted with ProLong™ Gold Antifade (Thermo Fisher Scientific) on Superfrost™ Plus microscope slides (Fisher Scientific), and fluorescent images were acquired with an Eclipse N*i* microscope (Nikon). Four fields of view per sample were analyzed to quantify percent of cells expressing MYOC, followed by calculation of mean ± SD as follows: DEX-induced MYOC upregulation = (MeanDEX – MeanCtrl) ± [(SDDEX)^2^ + (SDCtrl)^2^]^1/2^.

### Elastin-like polypeptide (ELP) expression and analysis

A plasmid containing an ELP sequence was kindly provided by Dr. Annabi (University of California, Los Angeles). It consists of 70 repeats of the pentapeptide VPGVG, in which the 1^st^ valine was replaced with isoleucine in every 5^th^ pentapeptide (i.e., ([VPGVG]4[IPGVG])14), flanked by KCTS residues to render the ELP UV-crosslinkable ^56^. *Escherichia coli (E. coli)* was used as a host to express the protein as described previously ^56,57^. Briefly, a 50 ml *E. coli* starter culture in terrific broth (Thermo Fisher Scientific) was incubated at 37°C overnight, followed by inoculation (1:100) of the expression culture and incubation at 37°C for 24 h. Bacterial pellets were collected and resuspended in lysis buffer (10 mM Tris, 1 mM EDTA, 100 mM NaCl, 5 mM MgCl2, 14.3 mM β-mercaptoethanol (BME); Thermo Fisher Scientific). Lysozyme (Acros Organics, Fair Lawn, NJ, USA) was added to the solution at 1 mg/ml and samples were incubated for 30 min on ice, followed by 5 min sonication (1 min on, 1 min off). The solution was centrifuged at 15,000 g for 20 min at 4°C and the supernatant was kept. To precipitate the protein, 1 M NaCl was added and centrifuged at 15,000 g for 20 min at 37°C. ELP was purified using inverse transition cycling (1 cold spin, one hot spin; x3), dialyzed against diH2O at 4°C for 18 h, and lyophilized. For characterization, ELP was reconstituted in DPBS at 10 mg/ml. Ten micrograms of ELP in 6X loading buffer (Boston Bio Products) with 5% BME (Fisher Scientific) were subjected to SDS-PAGE using NuPAGE™ 10% Bis-Tris Gels (Invitrogen) at 200V for 40 min. Gels were stained with Coomassie Blue R-250 (Biorad) and de-stained in a solution containing 50% (v/v) diH2O, 40% ethanol, and 10% acetic acid (Sigma-Aldrich). De-stained gels were imaged using a ChemiDoc system (BioRad) (**Suppl. Fig. S1**).

### Hydrogel precursor solutions

Methacrylate-conjugated bovine collagen type I (MA-COL; Advanced BioMatrix, Carlsbad, CA, USA) was reconstituted in sterile 20 mM acetic acid at 6 mg/ml. Immediately prior to use, 1 ml MA-COL was neutralized with 85 µl neutralization buffer (Advanced BioMatrix) according to the manufacturer’s instructions. Thiol-conjugated hyaluronic acid (SH-HA; Glycosil®; Advanced BioMatrix) was reconstituted in sterile diH2O containing 0.5% (w/v) photoinitiator (4-(2-hydroxyethoxy) phenyl-(2-propyl) ketone; Irgacure® 2959; Sigma-Aldrich) at 10 mg/ml according to the manufacturer’s protocol. In-house expressed ELP (SH-ELP; thiol via K<u>C</u>TS flanks) was reconstituted in DPBS at 10 mg/ml and sterilized using a 0.2 µm syringe filter in the cold.

### Preparation of PDMS molds

Custom master molds (8-mm diameter x 1-mm depth, or 8-mm diameter x 2-mm depth) were 3D-printed (F170; Stratasys, Eden Prairie, MN, USA) using ABS-M30 filament (Suppl. Fig. S2). Polydimethylsiloxane (PDMS; Sylgard 184, Dow Corning, Midland, MI, USA) was mixed in a 10:1 ratio of elastomer to curing agent according to the manufacturer’s protocol, poured into the 3D-printed negative master molds, degassed under vacuum in a desiccator, and cured overnight at 60°C. The PDMS molds were sterilized under UV light for 30 min prior to use.

### Preparation of HTM hydrogels

HTM cells (1.0×10^6^ cells/ml) were thoroughly mixed with MA-COL (3.6 mg/ml [all final concentrations]), SH-HA (0.5 mg/ml, 0.025% (w/v) photoinitiator), and SH-ELP (2.5 mg/ml) on ice. The HTM cell-laden hydrogel precursor solution was pipetted: a) onto PDMS-coated (Sylgard 184; Dow Corning) 24-well culture plates (10 µl droplets), b) into custom 8×1-mm PDMS molds (50 µl), c) onto 12-mm round glass coverslips (10 µl droplets) followed by spreading with pipette tips, or d) into standard 24-well culture plates (150 µl) depending on the type of experiment, and UV crosslinked (OmniCure S1500 UV Spot Curing System; Excelitas Technologies, Mississauga, Ontario, Canada) at 320-500 nm, 2.2 W/cm^2^ for 1-30 s before transferring constructs to 24-well culture plates (**Fig. 1**). One milliliter of DMEM with 10% FBS and 1% PSG was added to each well, and constructs were maintained at 37°C in a humidified atmosphere with 5% CO2. HTM hydrogels were cultured for 1-10 d with media replenished every 3 d.

**Fig. 1.**
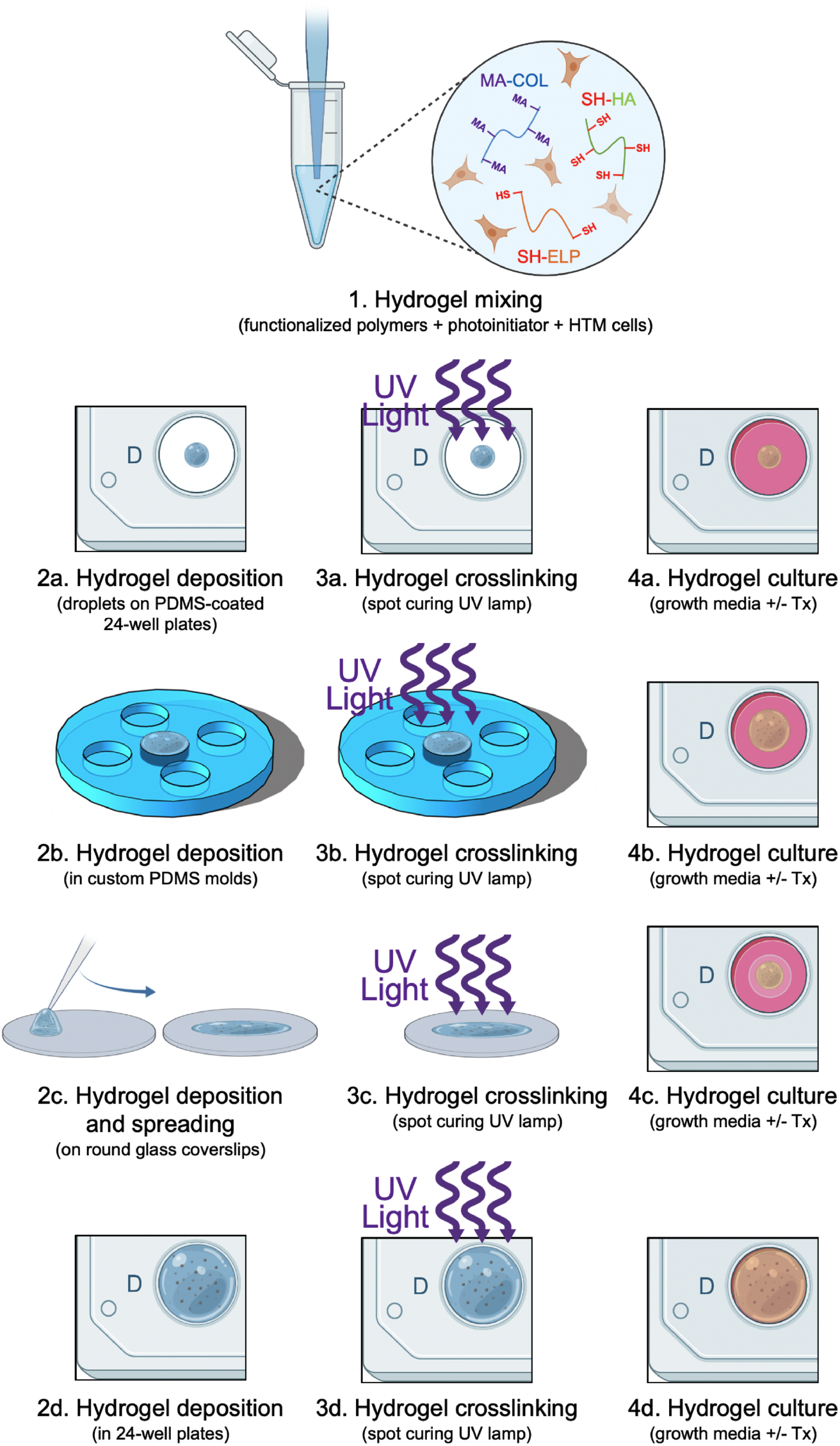
Schematic of HTM hydrogel formation. HTM hydrogels were fabricated by mixing HTM cells (1×10^6^ cells/ml) with methacrylate-conjugated collagen type I (MA-COL), thiol-conjugated hyaluronic acid (SH-HA; with photoinitiator), and in-house expressed elastin-like polypeptide (SH-ELP; thiol via K<u>C</u>TS flanks). Desired volumes of the HTM cell-laden hydrogel precursor solutions were added to different wells/molds [a) 10 µl on PDMS-coated 24-well plates, b) 50 µl in custom 8×1-mm PDMS molds, c) 10 µl on glass coverslips + spreading, d) 150 µl in standard 24-well culture plates], UV crosslinked (320-500 nm, 2.2 W/cm^2^, 1-30 s), and cultured in growth media, without or with different treatments, for 1-10 d.

### HTM hydrogel microstructural analysis

One hundred microliters of acellular hydrogel precursor solutions were pipetted into custom 8×2-mm PDMS molds, made from 3D-printed templates, UV crosslinked, and equilibrated in DPBS for 3 h to wash off any un-crosslinked polymers. Hydrogel samples were bisected to expose the interior hydrogel network surface and fixed in 4% PFA at 4°C overnight, washed with DPBS, flash frozen in liquid nitrogen, and lyophilized. Dried samples were mounted on scanning electron microscopy (SEM) stubs, sputter coated with gold, and visualized using a JSM-IT100 InTouchScope™ SEM (JEOL USA, Peabody, MA, USA) at 7 kV. Two fields of view per sample were analyzed to quantify hydrogel pore size (N = 40 total) using ImageJ software (National Institutes of Health, Bethesda, MD, USA).

### HTM hydrogel cell viability analysis

Cell viability was determined using a LIVE/DEAD™ Viability/Cytotoxicity Kit (i.e., live = green-stained, dead = red-stained) (Invitrogen) according to the manufacturer’s instructions. HTM hydrogels were incubated with the staining solutions (calcein-AM (0.5 μl/ml) and ethidium homodimer-1 (2 μl/ml) in DMEM with 10% FBS and 1% PSG) at 37°C for 45 min and washed with DPBS. Fluorescent images were acquired on 0 d (i.e., immediately after UV crosslinking; N = 4 per group and donor) and 7 d (qualitative) with an Eclipse T*i* microscope (Nikon). Four fields of view per sample were analyzed to quantify percent HTM cell viability (i.e., ratio of live to total cells), followed by calculation of mean ± SD.

### HTM hydrogel cell proliferation analysis

Cell proliferation was measured with the CellTiter 96® AQueous Non-Radioactive Cell Proliferation Assay (Promega, Madison, WI, USA) following the manufacturer’s protocol. HTM hydrogels cultured in DMEM with 10% FBS and 1% PSG for 1-10 d (N = 3 per group and donor) were incubated with the staining solution (38 μl MTS, 2 μl PMS solution, 200 μl DMEM) at 37°C for 1.5 h. Absorbance at 490 nm was recorded using a spectrophotometer plate reader (BioTEK, Winooski, VT, USA). Blank-subtracted absorbance values served as a direct measure of HTM cell proliferation over time.

### HTM hydrogel treatments

HTM hydrogels cultured in DMEM with 10% FBS and 1% PSG were subjected to the following treatments for 10 d: 1) vehicle control (ethanol; 0.1% (v/v)), 2) DEX (100 nM), 3) Rho-associated kinase (ROCK) inhibitor Y27632 (10 μM; Sigma-Aldrich), 4) DEX (100 nM) + Y27632 (10 μM) [co-treatment], or 5) DEX (100 nM) for 5 d followed by Y27632 (10 μM) for 5 d [sequential-treatment]. To validate our model, HTM hydrogels were subjected to: 1) vehicle control (4 mM hydrochloric acid with 0.1% bovine serum albumin (BSA); Fisher Scientific), 2) transforming growth factor beta-2 (TGF-β2; 2.5 ng/ml; R&D Systems, Minneapolis, MN, USA), 3) latrunculin-b (Lat-B; 2 μM; Fisher Scientific), 4) TGF-β2 (2.5 ng/ml) + Lat-B (2 μM) [co-treatment], or 5) TGF-β2 (2.5 ng/ml) for 5 d followed by Lat-B (2 μM) for 5 d [sequential-treatment].

### HTM hydrogel contraction analysis

HTM hydrogels were cultured in DMEM with 10% FBS and 1% PSG, in absence or presence of the different treatments (N = 3-4 per group and donor; 3 independent experiments). Longitudinal brightfield images were acquired over 10 d with an Eclipse T*i* microscope (Nikon). Construct area was measured using ImageJ software (NIH) over time and normalized to 0 d.

### HTM hydrogel immunocytochemistry analysis

HTM hydrogels, spread on glass coverslips and cultured in DMEM with 10% FBS and 1% PSG in presence of the different treatments for 10 d, were fixed with 4% PFA at 4°C overnight, permeabilized with 0.5% Triton™ X-100 (Thermo Fisher Scientific), blocked with blocking buffer (BioGeneX), and incubated with primary antibodies against α-smooth muscle actin (anti-αSMA [ab5694] 1:200; Abcam), followed by incubation with an Alexa Fluor® 488-conjugated secondary antibody; nuclei were counterstained with DAPI (both Abcam). Similarly, constructs were stained with Phalloidin-iFluor 488 (Abcam), nuclei were counterstained with DAPI, according to the manufacturer’s instructions. Coverslips were transferred to 35-mm glass bottom dishes (MatTek, Ashland, MA, USA), and fluorescent images were acquired with an Eclipse N*i* or T*i* microscope (Nikon).

### HTM hydrogel histology and immunohistochemistry analyses

HTM hydrogels, cultured in DMEM with 10% FBS and 1% PSG in presence of the different treatments for 10 d were fixed in 4% PFA at 4°C overnight, followed by incubation in 30% sucrose for 24 h at 4°C, washed with DPBS, embedded in Tissue-Plus™ O.C.T. Compound (Fisher Scientific), and flash frozen in liquid nitrogen. Twenty micrometer cryosections were cut using a cryostat (Leica Biosystems Inc., Buffalo Grove, IL, USA) and collected on Superfrost™ Plus microscope slides (Fisher Scientific). Sections were stained with picrosirius red/hematoxylin using the Picrosirius Red Stain Kit (PolySciences, Warrington, PA, USA) following the manufacturer’s protocol. Slides were mounted with Permount™ (Fisher Scientific) and brightfield images were captured with an Eclipse E400 microscope (Nikon). For immunohistochemistry analyses, sections were permeabilized with 0.5% Triton™ X-100, blocked with blocking buffer, and incubated with primary antibodies against fibronectin (anti-fibronectin [ab45688] 1:500; Abcam) or collagen type IV (anti-collagen IV [ab6586] 1:500; Abcam) followed by incubation with an Alexa Fluor® 488-conjugated secondary antibody; nuclei were counterstained with DAPI (both Abcam). Slides were mounted with ProLong™ Gold Antifade (Thermo Fisher Scientific), and fluorescent images were acquired with an Eclipse N*i* microscope (Nikon). Three fields of view per sample were analyzed to quantify normalized fibronectin signal intensity, followed by calculation of fold-change vs. control.

### HTM hydrogel rheology analysis

Fifty microliters of HTM cell-laden or acellular hydrogel precursor solutions were pipetted into custom 8×1-mm PDMS molds (**Suppl. Fig. S2**). Similarly, 150 µl of HTM cell-laden hydrogel precursor solutions were pipetted into 24-well culture plates. All samples were UV crosslinked and equilibrated as described above. Acellular hydrogels were measured on 0 d. HTM hydrogels, cultured in DMEM with 10% FBS and 1% PSG in presence of the different treatments, were measured on 0 d and 10 d; plates were scanned for visualization followed by cutting samples to size using an 8-mm diameter tissue punch. A Kinexus rheometer (Malvern Panalytical, Westborough, MA, USA) fitted with an 8-mm diameter parallel plate was used to measure hydrogel viscoelasticity. To ensure standard conditions across all experiments (N = 3 per group), the geometry was lowered into the hydrogels until a calibration normal force of 0.02 N was achieved. Subsequently, an oscillatory shear-strain sweep test (0.1-60%, 1.0 Hz, 25°C) was applied to determine storage modulus (G’) and loss modulus (G”) in the linear region. Compression testing was performed at axial strains from 0% to −50% by changing the gap between the plates followed by applying a sinusoidal shear strain of 2% at a frequency of 1.0 Hz to determine storage modulus (G’) with increasing axial strain.

### Statistical analysis

Individual sample sizes and details of statistical analyses are specified in each figure caption. Comparisons between groups were assessed by unpaired *t* test, one-way or two-way analysis of variance (ANOVA) with Tukey’s multiple comparisons *post hoc* tests, as appropriate. All data are shown with mean ± SD, some with individual data points. The significance level was set at p<0.05 or lower. GraphPad Prism software v8.4 (GraphPad Software, La Jolla, CA, USA) was used for all analyses.

## Results

### HTM cell characterization

A reliable feature of HTM cells *in vitro* is upregulation of MYOC expression in more than 50% of cells in response to challenge with the corticosteroid DEX ^54^. Three strains (HTM01, HTM11, HTM12) were used and compared to a validated reference strain (HTM129). Both HTM01 and HTM12 exhibited normal morphology (i.e., cobblestone-like pattern with some overlapping processes; **Fig. 2A**) and growth characteristics (i.e., contact inhibited with doubling time of ~2 d) comparable to the reference standard. HTM11 showed overall normal morphology; however, a fraction of cells was noticeably larger. An overall slower doubling time (~4 d) was also noted. Significantly increased MYOC mRNA expression by qRT-PCR was observed with DEX treatment vs. controls across HTM cell strains (**Fig. 2B**). Qualitative assessment of secreted MYOC protein by immunoblot showed robust DEX-inducible expression (**Fig. 2C**). Intracellular MYOC expression by immunocytochemistry was increased by 57.9 ± 7.2% (reference), 81.5 ± 1.9% (HTM01), 41.5 ± 11.4% (HTM11), and 51.8 ± 6.6% (HTM12) with DEX treatment vs. controls (p<0.001 for all HTM cell strains; **Fig. 2D**). Alpha B-Crystallin (CRYAB) is exclusively expressed in the JCT region of the HTM ^58^. Collagenase was used to digest the dissected strips of HTM tissue; specifically, to disrupt contacts between JCT-HTM cells and their dense ECM, and to encourage cell migration out of the HTM tissue. Our results showed that all of our HTM cell strains and the reference cells highly expressed CRYAB (**Fig. 2E**), suggesting that a significant population of cells were derived from the JCT layer. Based on suboptimal cellular morphology and growth characteristics, as well as DEX-inducible intracellular MYOC expression below the threshold of 50%, HTM11 cells were ruled out for the rest of the studies.

**Fig. 2.**
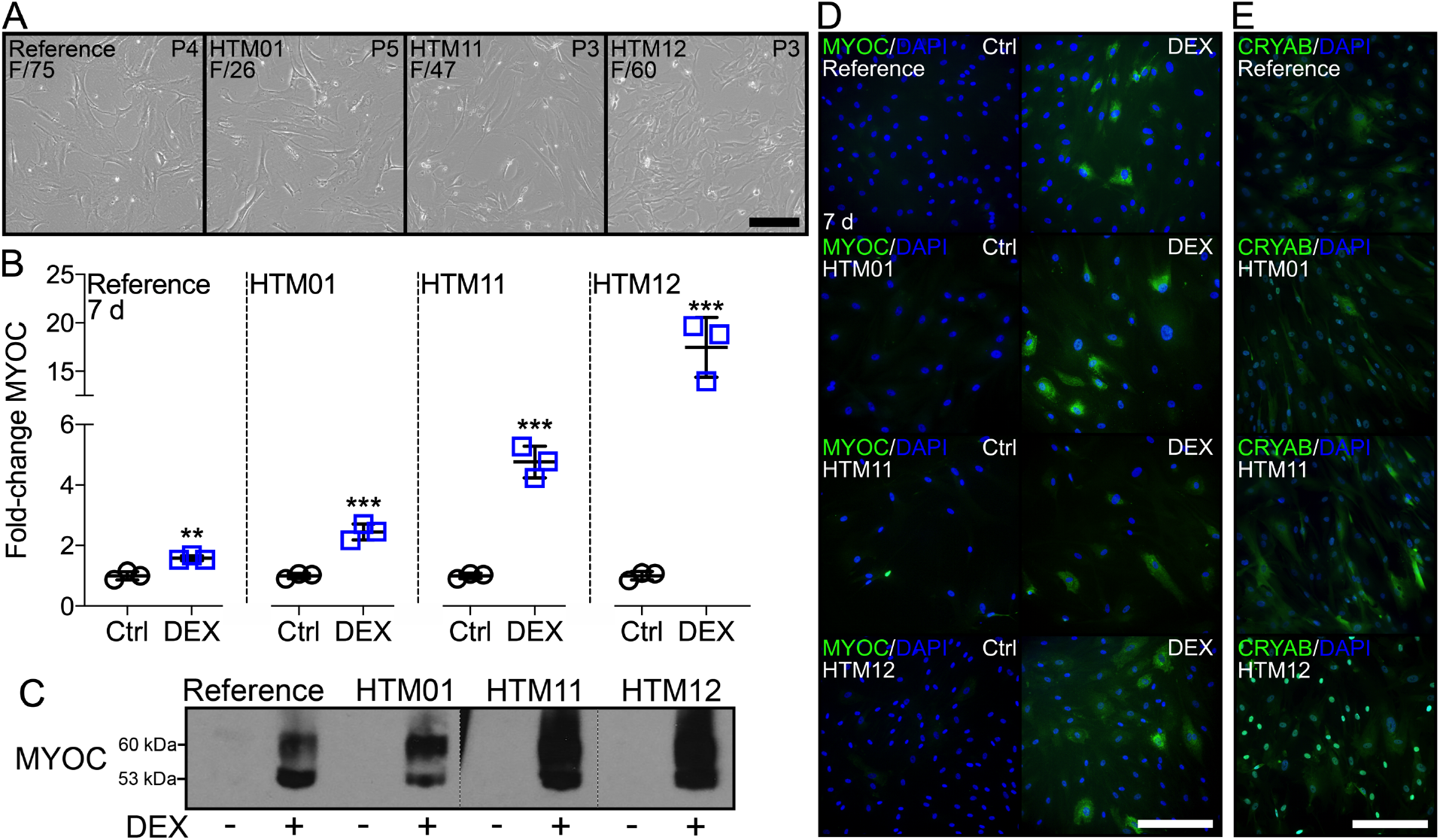
HTM cell characterization. (**A**) Representative phase contrast micrographs of reference, HTM01, HTM11, and HTM12 cell strains with sex/age information. Scale bar, 250 µm. (**B**) mRNA fold-change of MYOC by qRT-PCR at 7 d (N = 3 per group and donor; *p<0.05, ***p<0.001 vs. control). (**C**) Immunoblot of secreted MYOC at 7 d. (**D**) Representative fluorescence micrographs of intracellular MYOC at 7 d (MYOC = green; DAPI = blue). Scale bar, 250 μm. (**E**) Representative fluorescence micrographs of CRYAB (CRYAB = green; DAPI = blue). Scale bar, 250 μm.

Together, this data suggests that HTM01 and HTM12 exhibit all required key characteristics according to a recent consensus paper ^54^ to faithfully identify them as normal HTM cells, comparable to a confirmed reference standard.

### HTM cell viability in hydrogels

The number of HTM cells in native tissue decreases with aging. A steady decline from (0.59-1.74)x10^6^ cells/ml in 20-year-old individuals to (0.31-0.92)x10^6^ cells/ml in 80-year-old individuals has been shown ^7,59^. Therefore, we used a normalized density of 1.0×10^6^ cells/ml in HTM cell-laden hydrogels independent of donor age. To ascertain that HTM cells could withstand the UV light-activated free-radical crosslinking process, referred to as photocrosslinking or UV crosslinking throughout, HTM cell viability in hydrogels crosslinked for increasing times was determined by live/dead staining. Hydrogel-encapsulated HTM cells displayed high viability immediately after photocrosslinking (0 d). We observed negligible cell death across HTM cell strains after 1 s and 5 s of UV crosslinking, and more than 80% cells remained viable after 10 s and 20 s of irradiation. In contrast, significant cell death was noted after 30 s of UV crosslinking (**Fig. 3**).

**Fig. 3.**
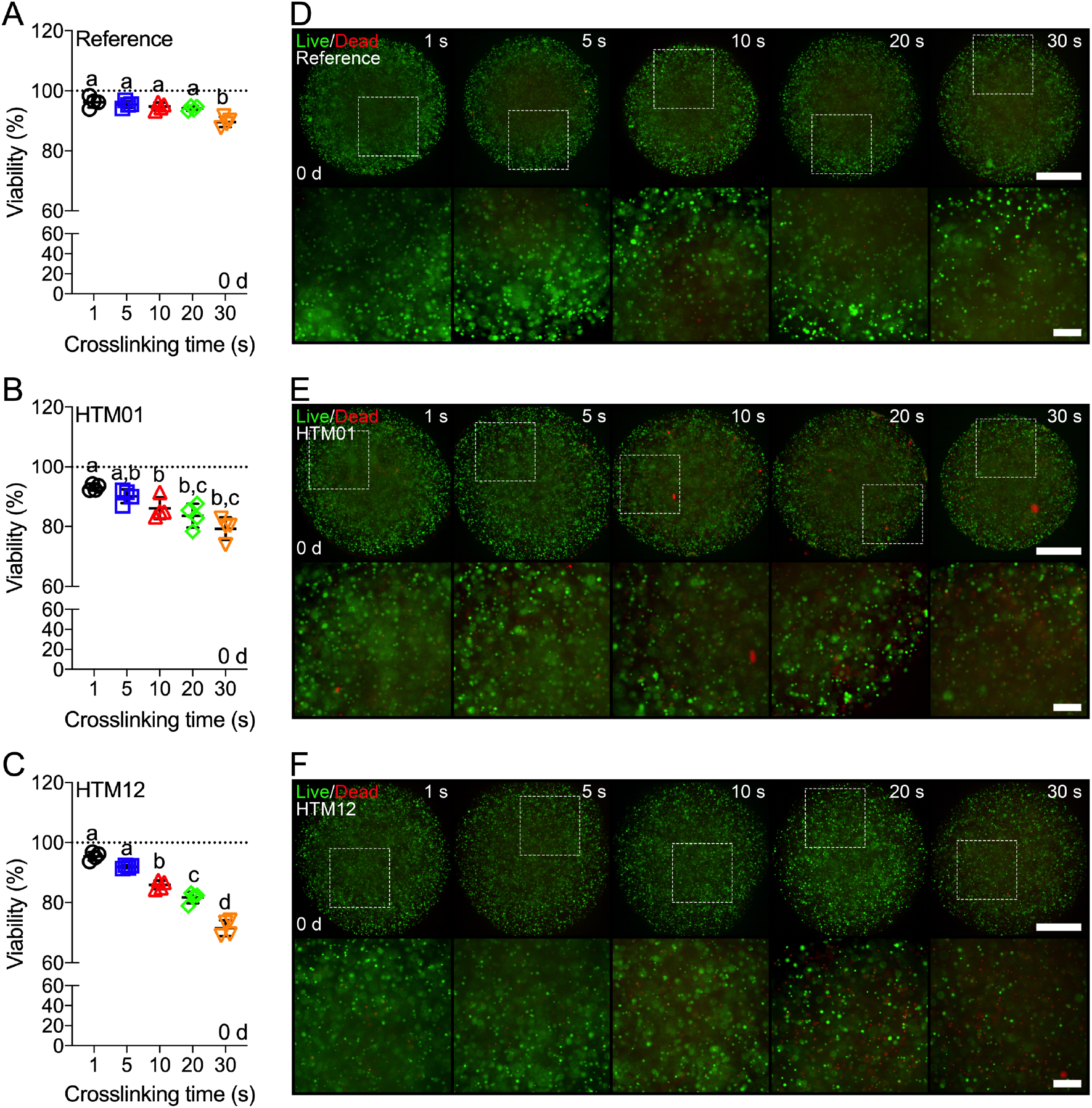
HTM cell viability. Cell viability quantification of hydrogel-encapsulated (**A**) reference, (**B**) HTM01, and (**C**) HTM12 cells across UV crosslinking times at 0 d (N=4 per group and donor; dotted lines show 100% viability for reference; shared significance indicator letters represent non-significant difference (p>0.05), distinct letters represent significant difference (p<0.05)). Representative fluorescence micrographs of (**D)** reference, (**E**) HTM01, and (**F**) HTM12 hydrogels at 0 d (live cells = green; dead cells = red). Scale bars, 1 mm (top) and 250 μm (bottom).

This suggests that UV irradiation for 1-10 s does not cause major cytotoxicity to HTM cells encapsulated in 3D biopolymer hydrogels, regardless of donor age and cell source. Therefore, we focused on HTM01 and HTM12 isolated in our laboratory for subsequent experiments.

### HTM cell contractility and proliferation in hydrogels

HTM cells interact with and contract their ECM environment ^10^. For both HTM01 and HTM12 cell strains, an inverse relationship between UV crosslinking time and hydrogel contraction was observed; i.e., constructs showed overall decreased contraction with increasing UV crosslinking time (**Fig. 4; Suppl. Fig. S3**). This was consistent with hydrogel stiffness; i.e., acellular constructs exhibited increased shear storage moduli (G’) by rheology with increasing UV crosslinking time, while loss moduli (G”) remained relatively stable (**Suppl. Fig. S4**). A key advantage of using hydrogel microenvironments to assess cellular behaviors vs. traditional 2D or scaffold-based 3D culture systems is their viscoelastic behavior ^50^. In this study, hydrogels showed tissue-like viscoelastic properties independent of UV crosslinking time, with significantly higher G’ vs. G” indicating predominantly elastic rather than viscous characteristics (**Suppl. Fig. S4**).

**Fig. 4.**
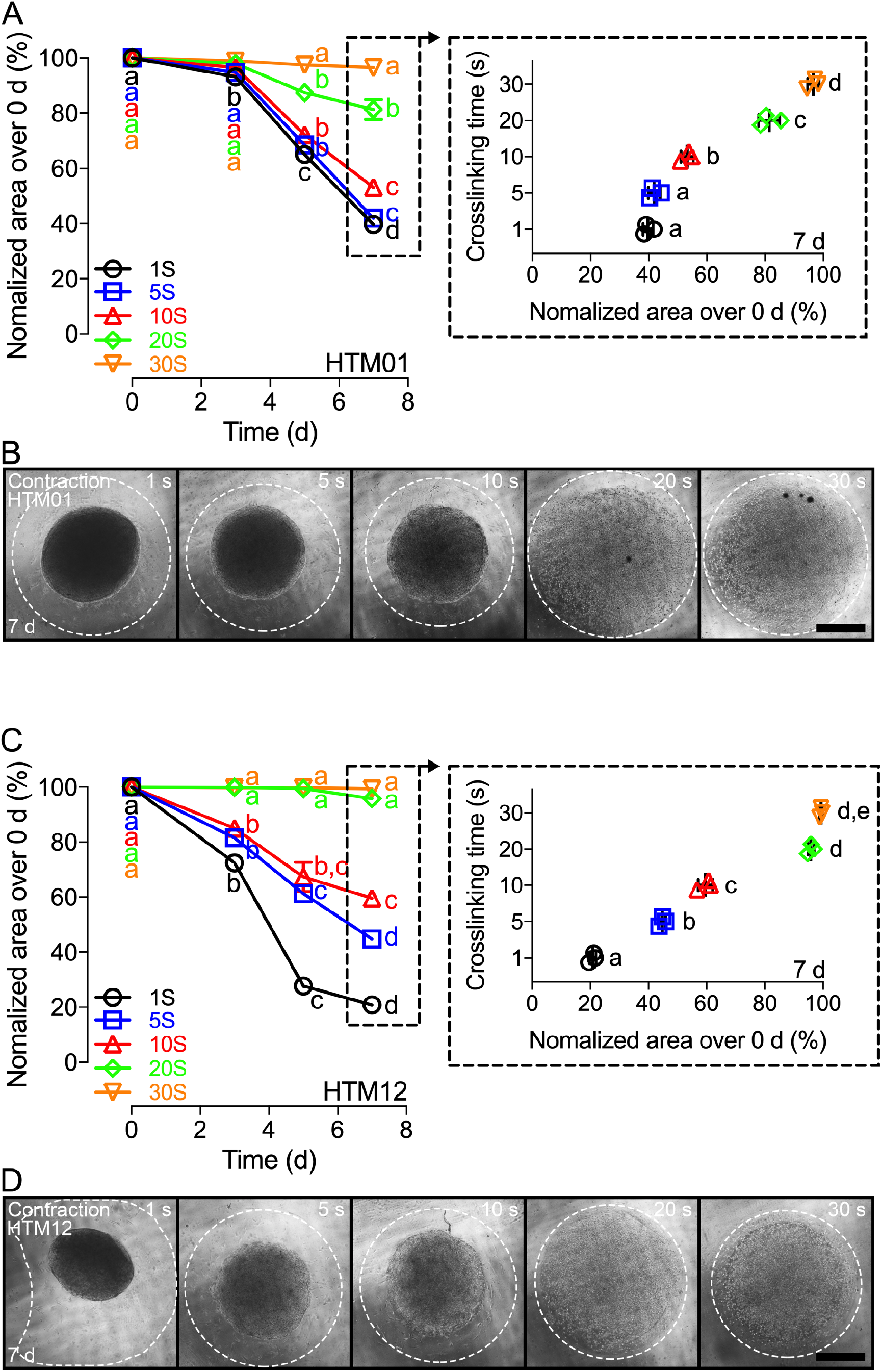
HTM hydrogel contractility. Longitudinal quantification (i.e., construct size relative to 0 d) of (**A**) HTM01 and (**C**) HTM12 cell contractility in hydrogels across crosslinking times; dashed boxes show detailed comparisons between groups at 7 d (N=3 per group and donor; shared significance indicator letters represent non-significant difference (p>0.05), distinct letters represent significant difference (p<0.05)). Representative brightfield images of (**B)** HTM01 and (**D**) HTM12 hydrogels at 7 d; white dashed lines outline original size of constructs at 0 d. Scale bars, 1 mm.

HTM01 hydrogels crosslinked for 1-10 s significantly contracted over culture time, reaching ~40-50% of their original size by 7 d. No differences between constructs irradiated for 1 s and 5 s were found, but hydrogels formed after 10 s UV crosslinking were significantly less contracted in comparison (**Fig. 4A,B; Suppl. Fig. S3A**). Constructs in the 20 s group also displayed significant longitudinal contraction, approximating ~80% of their baseline size by 7 d at which point they were significantly less contracted vs. all shorter UV crosslinking groups. HTM01 hydrogels crosslinked for 30 s displayed consistent size over time, and were significantly less contracted vs. all other groups at 7 d (**Fig. 4A,B; Suppl. Fig. S3A**). HTM12 hydrogels showed similar overall behavior to HTM01 (**Fig. 4C,D; Suppl. Fig. S3B**). Of note, constructs UV crosslinked for 1 s were significantly more contracted compared to all other groups by 7 d, reaching ~20% of their original size. No differences between constructs irradiated for 20 s and 30 s were found; hydrogels were significantly less contracted vs. all shorter UV crosslinking groups (**Fig. 4C,D; Suppl. Fig. S3B**). This data suggests that HTM cells are able to contract and deform the hydrogels photocrosslinked for 1-10 s independent of cell strain.

Taken together, consistent hydrogel formation was observed with 5 s UV crosslinking. Using these conditions, constructs showed high HTM cell viability immediately after hydrogel formation (**Fig. 3**) together with reliable contractile-mediated hydrogel deformation over time (**Fig. 4; Suppl. Fig. S3**). This was accompanied by significant HTM cell proliferation in hydrogels over 7 d (**Fig. 5A,B**), at which point negligible cell death and prominent cell spreading were observed with both HTM cell strains (**Fig. 5C**). Therefore, 5 s UV crosslinking time was chosen for all following experiments with focus on HTM12.

**Fig. 5.**
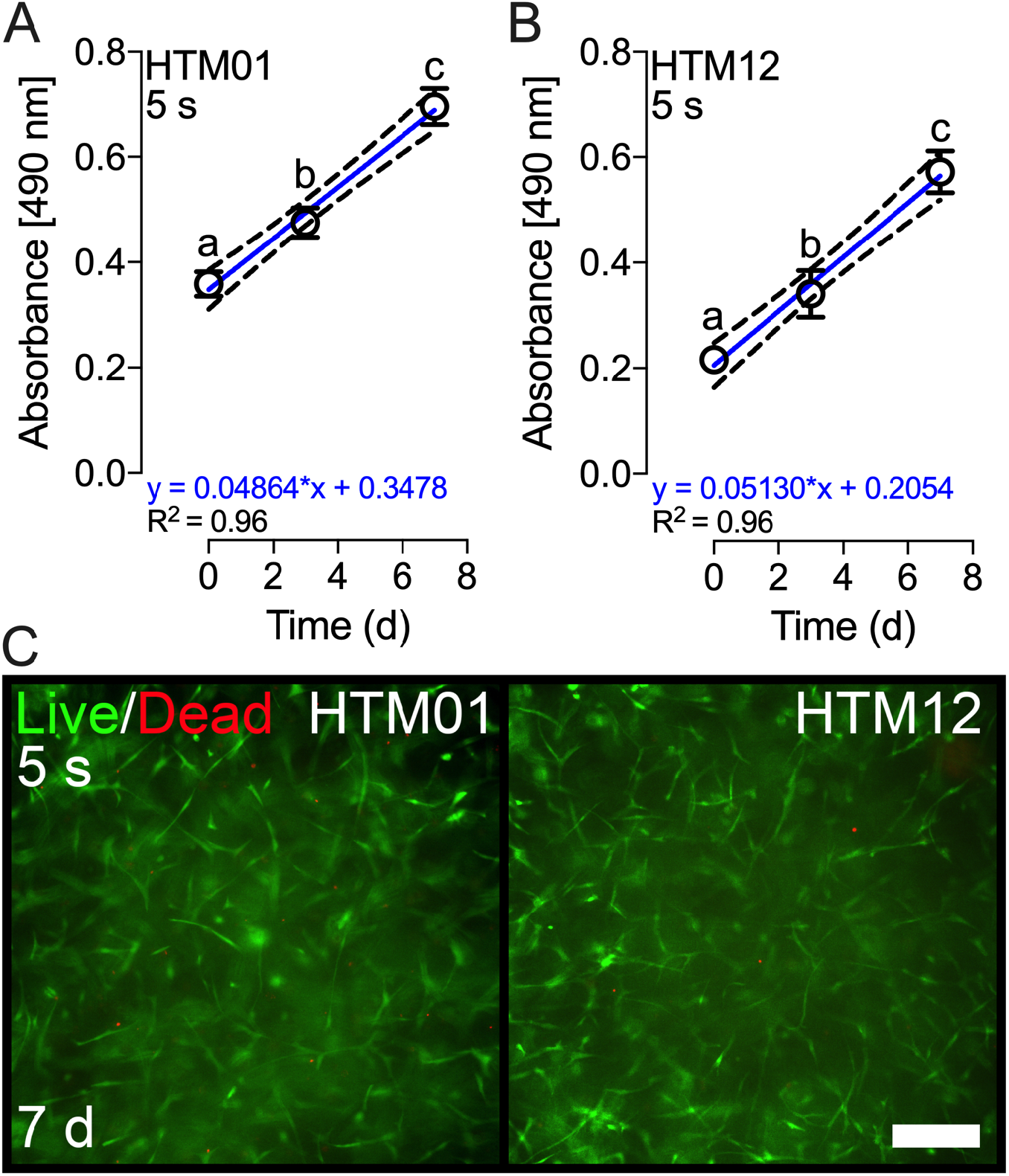
HTM cell proliferation and viability in hydrogels. Longitudinal cell proliferation quantification of (**A**) HTM01 and (**B**) HTM12 hydrogels crosslinked for 5 s (N=3 per group and donor; shared significance indicator letters represent non-significant difference (p>0.05), distinct letters represent significant difference (p<0.05)). (**C**) Representative fluorescence micrographs of HTM01 and HTM12 hydrogels crosslinked for 5 s at 7 d (live cells = green; dead cells = red). Scale bar, 250 μm.

### HTM hydrogel microstructural and mechanical properties

HTM cells within the JCT region reside in a loose connective tissue comprised of reticular and elastic fibers, and ground substance made of proteoglycans and glycosaminoglycans. Pore sizes across the JCT have been estimated at ~2-15 µm ^3–7^. To assess the hydrogel microarchitecture, SEM images were acquired on 0 d (i.e., immediately after 5 s UV crosslinking). Acellular hydrogels exhibited a honeycomb-like architecture with 43.3 ± 27.3 µm pore size (**Fig. 6A**), comparable to a recently reported hydrogel made of ELP and HA ^57^. For correlative analyses, we next assessed hydrogel mechanical properties by rheology of both acellular and HTM12 cell-encapsulated hydrogels. In general, hydrogel stiffness is dependent on biopolymer concentrations/arrangement and crosslinking density, with encapsulated contractile cells further contributing to construct elasticity. At the chosen crosslinking time of 5 s, we observed overall relatively low modulus values in the low pascal-range, consistent with the low-to-medium biopolymer concentrations used. HTM cell-encapsulated hydrogels were significantly stiffer compared to acellular samples (G’, ~75 Pa vs. 47 Pa), while G” was comparable (**Fig. 6B**). Regardless of whether or not cells were present within the 3D biopolymer network, hydrogels displayed tissue-like compression stiffening ^60^ (i.e., increase in shear modulus with increasing compressive strain) (**Fig. 6C**).

**Fig. 6.**
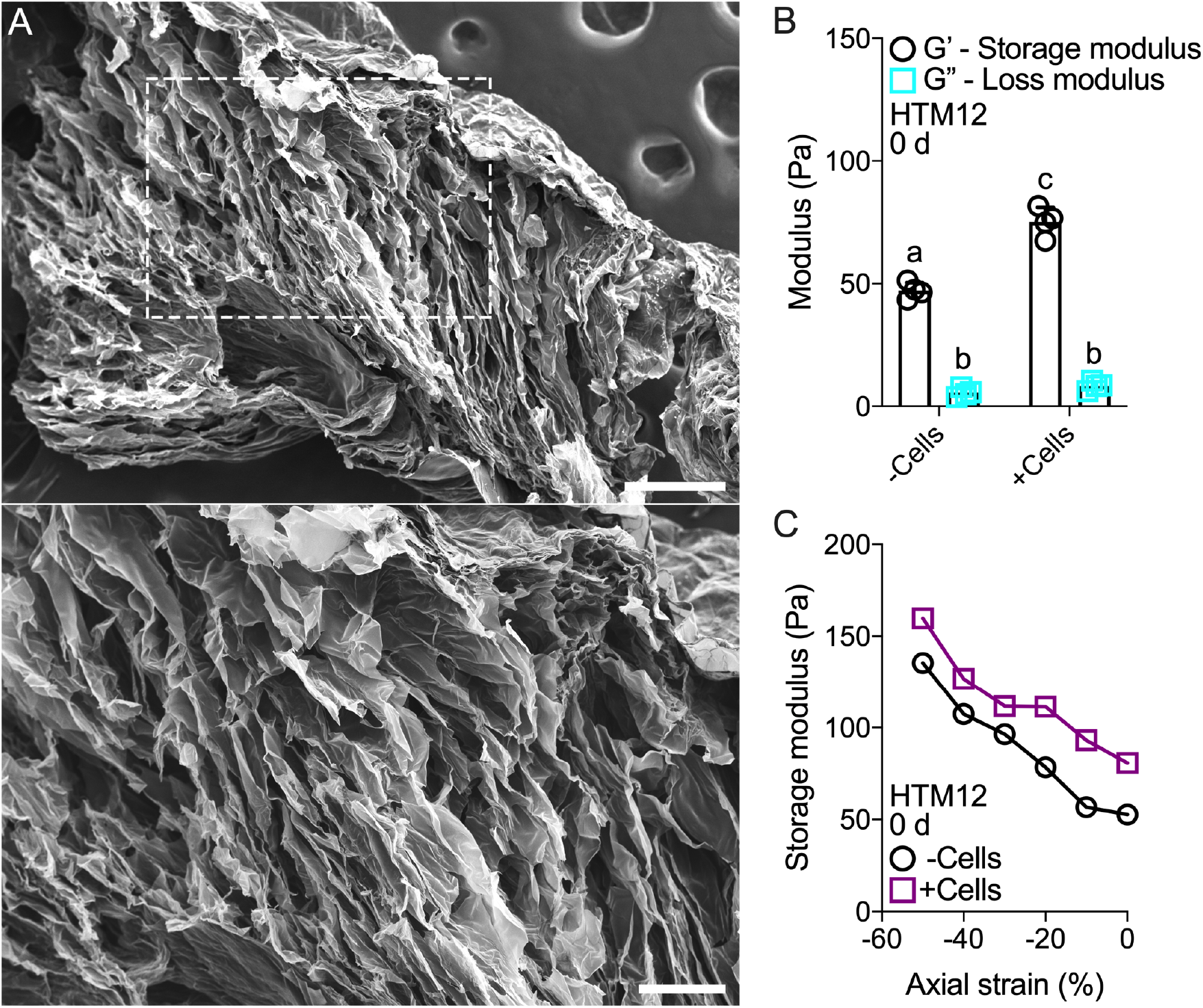
HTM hydrogel microstructural and mechanical analyses. (**A**) Scanning electron micrographs of acellular hydrogels crosslinked for 5 s at 0 d (dashed box shows region of interest in higher magnification image). Scale bars, 250 µm (top) and 100 µm (bottom). (**B**) Storage and loss moduli of acellular and HTM12 cell-encapsulated hydrogels crosslinked for 5 s at 0 d (N=4 per group; shared significance indicator letters represent non-significant difference (p>0.05), distinct letters represent significant difference (p<0.05)). (**C**) Storage modulus measured at 2% shear strain as a function of axial strain for acellular and HTM12 cell-encapsulated hydrogels crosslinked for 5 s at 0 d (N=1 per group).

This data suggests that our HTM hydrogel system provides an adequately porous and soft 3D microenvironment that mimics the mechanical properties of native heterogeneous soft connective tissues.

### HTM cell-laden hydrogels as glaucoma disease model: pharmacological induction/rescue

#### Hydrogel contractility

The contractility status of HTM cells and overall tissue influences AH outflow resistance and IOP ^61^. Increased HTM contraction is observed in patients with POAG, which typically leads to tissue stiffening, decreased AH outflow facility, and elevated IOP ^59^. Steroid therapy is widely used to treat a variety of inflammatory diseases and conditions. However, long term use of ocular corticosteroids such as DEX often results in elevated IOP and POAG ^62^. To ascertain whether photocrosslinked HTM hydrogels could serve as a glaucoma disease model, we treated constructs with DEX ^54^ to induce POAG-like conditions and evaluated its effects on hydrogel contractility. In the United States, the ROCK inhibitor netarsudil is a clinically available drug that directly targets the stiffened HTM to increase AH outflow via cell/tissue relaxation ^21–25^. Therefore, we assessed the therapeutic effects of ROCK inhibitor Y27632 treatment on HTM hydrogel contractility following DEX induction.

We observed significant contraction over time in all groups (**Fig. 7A; Suppl. Fig. S5**). DEX-treated HTM hydrogels exhibited significantly greater contraction vs. controls by 5 d, reaching ~44% of their original size. Comparable values were found in the sequential treatment samples (DEX (5 d) + Y27632 (5 d)), which at this point had only been exposed to DEX rendering the two groups virtually identical. In contrast, Y27632 significantly relaxed the HTM hydrogels (i.e., decreased contraction; ~70% of their original size) vs. controls. No differences were noted between DEX + Y27632 co-treated samples and controls by 5 d, displaying significantly less contraction compared to DEX- and DEX (5 d) + Y27632 (5 d)-treated HTM hydrogels, yet significantly more contraction vs. Y27632-treated constructs; this suggests that Y27632 can prevent DEX-induced pathologic contraction of HTM hydrogels when presented together (**Fig. 7A,B,D; Suppl. Fig. S5**).

**Fig. 7.**
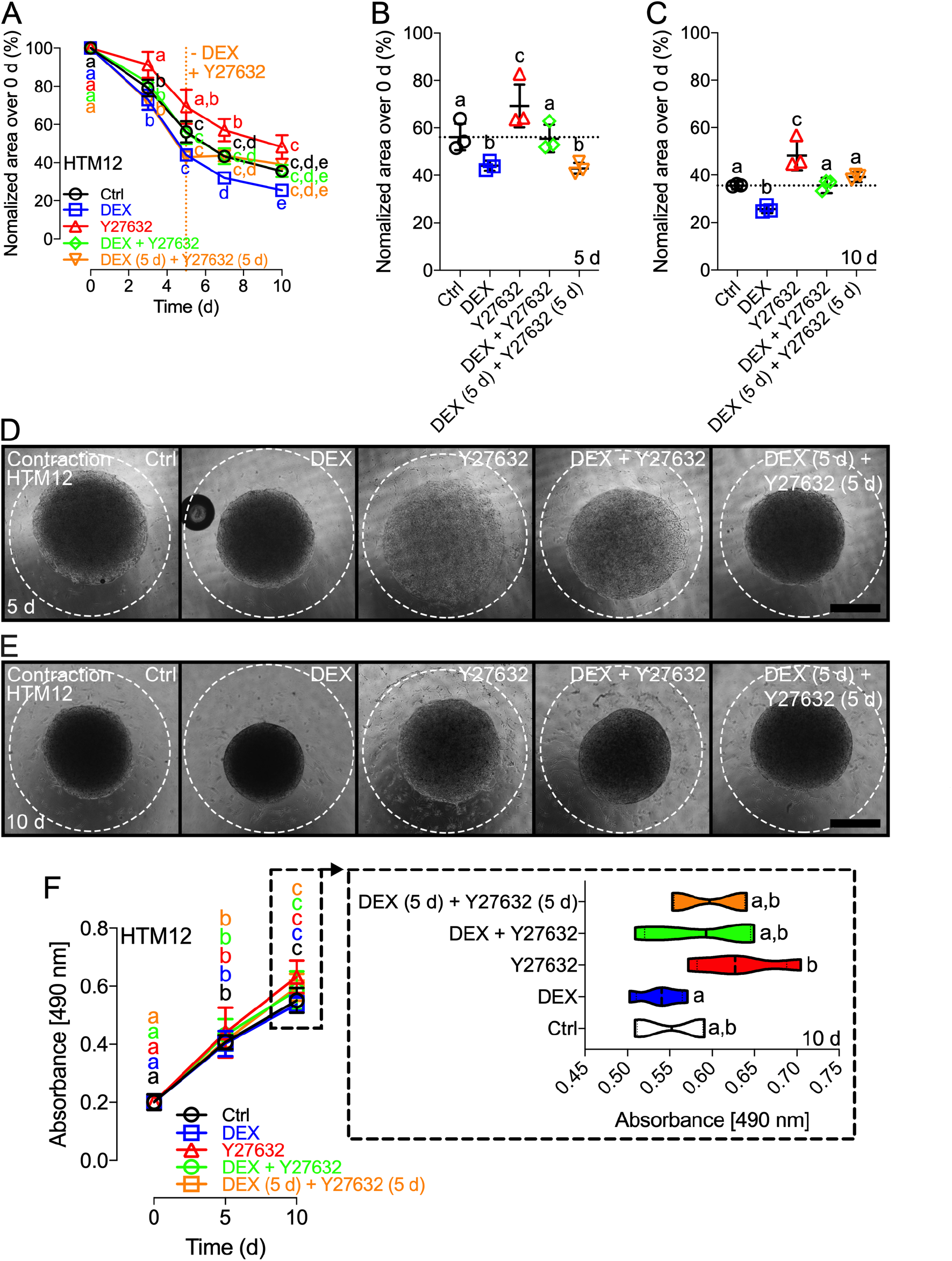
HTM hydrogel contractility and cell proliferation with corticosteroid induction and ROCK inhibitor rescue. (**A**) Longitudinal quantification (i.e., construct size relative to 0 d) of HTM12 cell contractility in hydrogels subjected to control, 100 nM DEX, 10 µM Y27632, DEX + Y27632, or DEX (5 d) + Y27632 (5 d) (N=3 per group; shared significance indicator letters represent non-significant difference (p>0.05), distinct letters represent significant difference (p<0.05)). Detailed comparisons between groups at (**B**) 5 d and (**C**) 10 d (dotted lines show respective control values for reference). Representative brightfield images of HTM12 hydrogels subjected to the different treatments at (**D**) 5 d and (**E**) 10 d; white dashed lines outline original size of constructs at 0 d. Scale bars, 1 mm. (**F**) Longitudinal cell proliferation quantification of HTM12 hydrogels subjected to the different treatments; dashed box shows detailed comparisons between groups at 10 d (N=3 per group; shared significance indicator letters represent non-significant difference (p>0.05), distinct letters represent significant difference (p<0.05)).

Overall similar trends were observed at 10 d (**Fig. 7A,C,E; Suppl. Fig. S5**). DEX treatment induced significantly more contraction (~26% of their original size), whereas Y27632 treatment resulted in significantly less contraction (~48% of their original size) vs. all other groups. No differences were observed between DEX + Y27632 co-treatment and controls. Importantly, in a clinical scenario, the HTM tissue would likely already show signs of dysfunction before any POAG treatment would be administered. We simulated this with the sequential treatment group: DEX induction for 5 d followed by Y27632 treatment for 5 d with DEX withheld. Our results showed that Y27632 rescued HTM hydrogel contraction induced by DEX and prevented further contraction; i.e., construct size with DEX (5 d) + Y27632 (5 d) sequential treatment was similar to the control and DEX + Y27632 groups, but significantly increased compared to DEX-treated samples (**Fig. 7A,C,E; Suppl. Fig. S5**).

Next, to see if hydrogel contractility was influenced by the cell number, we assessed HTM cell proliferation in constructs subjected to the different treatments. Significant HTM cell proliferation over time was observed in all groups (**Fig. 7F**). No differences between groups were found at 5 d; however, by 10 d HTM cells in the DEX-treated group showed significantly less proliferation compared to the Y27632 treated group (**Fig. 7F; inset in dashed box**).

Together, this suggests that DEX robustly induces HTM hydrogel contractility mimicking glaucomatous conditions, and that ROCK inhibition prevents or rescues DEX-induced pathologic contraction.

To validate the utility of the HTM hydrogel as glaucoma disease model, we next assessed an alternative pair of pharmacological induction and rescue: transforming growth factor beta-2 (TGF-β2), implicated in the pathogenesis of POAG ^63^, and latrunculin-b (Lat-B), which prevents polymerization of actin filaments ^64^. Remarkably similar results to DEX/Y27632 were obtained. We noted significant contraction over time in all groups (**Suppl. Fig. S6A**). TGF-β2-treated HTM hydrogels showed significantly greater contraction vs. controls by 5 d, reaching ~45% of their original size. Comparable values were observed in the sequential treatment samples (TGF-β2 (5 d) + Lat-B (5 d); identical to TGF-β2 at this time point). Lat-B only slightly relaxed the HTM hydrogels (~58% of their original size) vs. controls. Furthermore, no differences were found between TGF-β2 + Lat-B co-treated samples and controls by 5 d, but constructs were significantly less contracted compared to TGF-β2-treated HTM hydrogels; this suggests that Lat-B can offset TGF-β2-induced pathologic contraction of HTM hydrogels when presented together (**Suppl. Fig. S6A,B,D**). By 10 d, TGF-β2 treatment induced significantly more contraction (~25% of their original size) vs. all other groups, while Lat-B treatment resulted in only slightly less contraction (~44% of their original size) vs. controls. No differences were observed between TGF-β2 + Lat-B co-treatment and controls. However, Lat-B rescued HTM hydrogel contraction induced by TGF-β2; i.e., construct size with TGF-β2 (5 d) + Lat-B (5 d) sequential treatment was similar to the control and TGF-β2 + Lat-B groups, but significantly increased compared to TGF-β2-treated samples (**Suppl. Fig. S6A,C,E**).

### Hydrogel actin rearrangement

The actin cytoskeleton, the primary force-generating machinery in cells, plays fundamental roles in various cellular process such as migration, morphogenesis, and cytokinesis ^65^. <u>F</u>ilamentous f-actin fiber arrangement directly affects cell and tissue contraction. Therefore, we next investigated f-actin and alpha smooth muscle actin (αSMA) abundance and organization in HTM hydrogels subjected to the different treatments (**Fig. 8**). DEX-treated constructs showed substantial reorganization of HTM cell microfilaments, with increased formation of f-actin/stress fibers, minor actin tangles, and crosslinked actin network (CLAN)-like structures in the 3D hydrogel environment vs. controls. By contrast, Y27632 treatment depolarized microfilaments with notably decreased f-actin fibers compared to both control and DEX-treated groups. No differences were observed for DEX + Y27632 co- or sequential treatment in comparison to Y27632; again markedly decreased f-actin fibers were noted vs. control and DEX-treated groups (**Fig. 8A**). Overall relatively similar trends were observed for αSMA. DEX-treated samples showed substantially increased αSMA expression vs. all other groups, whereas Y27632 nearly abolished the αSMA signal (high exposure chosen to illustrate this point). Again, no differences were observed between DEX + Y27632 co- or sequential treatment, with comparable αSMA expression to controls, but higher levels vs. Y27632-treated hydrogels (**Fig. 8B**).

**Fig. 8.**
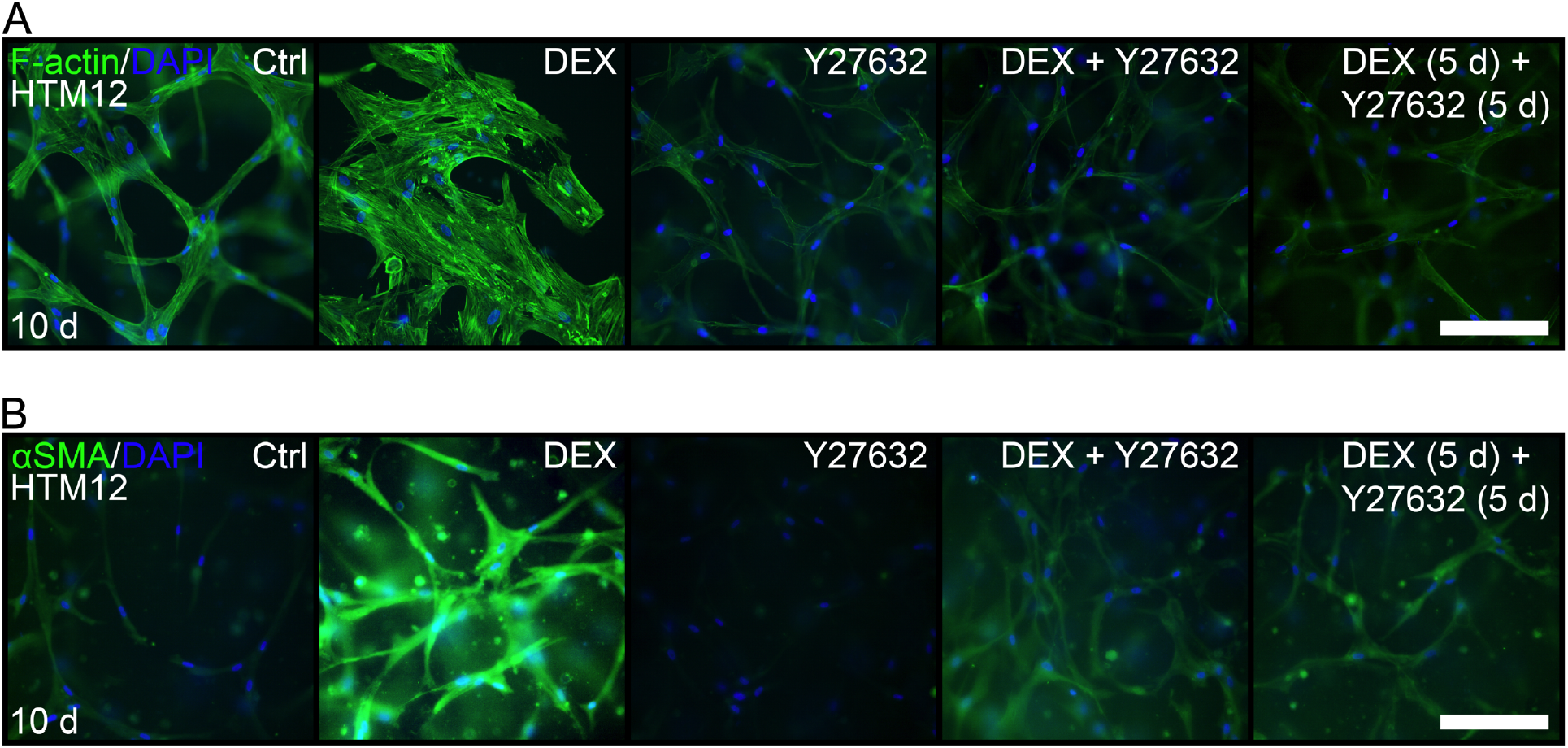
F-actin and αSMA expression by HTM cells in hydrogels with corticosteroid induction and ROCK inhibitor rescue. Representative fluorescence micrographs of (**A**) f-actin and (**B**) αSMA in HTM12 hydrogels subjected to control, 100 nM DEX, 10 µM Y27632, DEX + Y27632, or DEX (5 d) + Y27632 (5 d) at 10 d (f-actin and αSMA = green; DAPI = blue). Scale bars, 250 μm.

Together, these data show that DEX increases f-actin fibers and α-SMA in HTM hydrogels, which is potently rescued by Y27632. This is consistent with the reported reduction of actomyosin contractile tone in HTM tissue following ROCK inhibition ^22–25^. These changes in f-actin and α-SMA could have a direct impact on HTM cell contractility and interactions with their surrounding ECM, as well as HTM tissue stiffness and ultimately AH outflow resistance.

### Hydrogel morphology and ECM deposition

Continuous remodeling of baseline hydrogel ECM components (i.e., fibrillar collagen type I and ELP, plus HA) and HTM cell-secreted ECM over culture time, and in response to the different treatments is expected and desired. We next investigated overall morphology of HTM hydrogels subjected to the different treatments using picrosirius red-stained sections (**Fig. 9A**). Consistent with the contractility analyses, DEX-treated HTM hydrogels appeared condensed with decreased inter-fibrillar spaces and reduced fiber thickness vs. controls. By contrast, Y27632 noticeably increased the inter-fibrillar spaces compared to controls, while fiber thickness appeared largely unaffected. DEX + Y27632 co- or sequential treatment were comparable to controls but overall less dense vs. DEX-treated constructs. Of note, abundant hematoxylin-stained HTM cells were visible across groups.

**Fig. 9.**
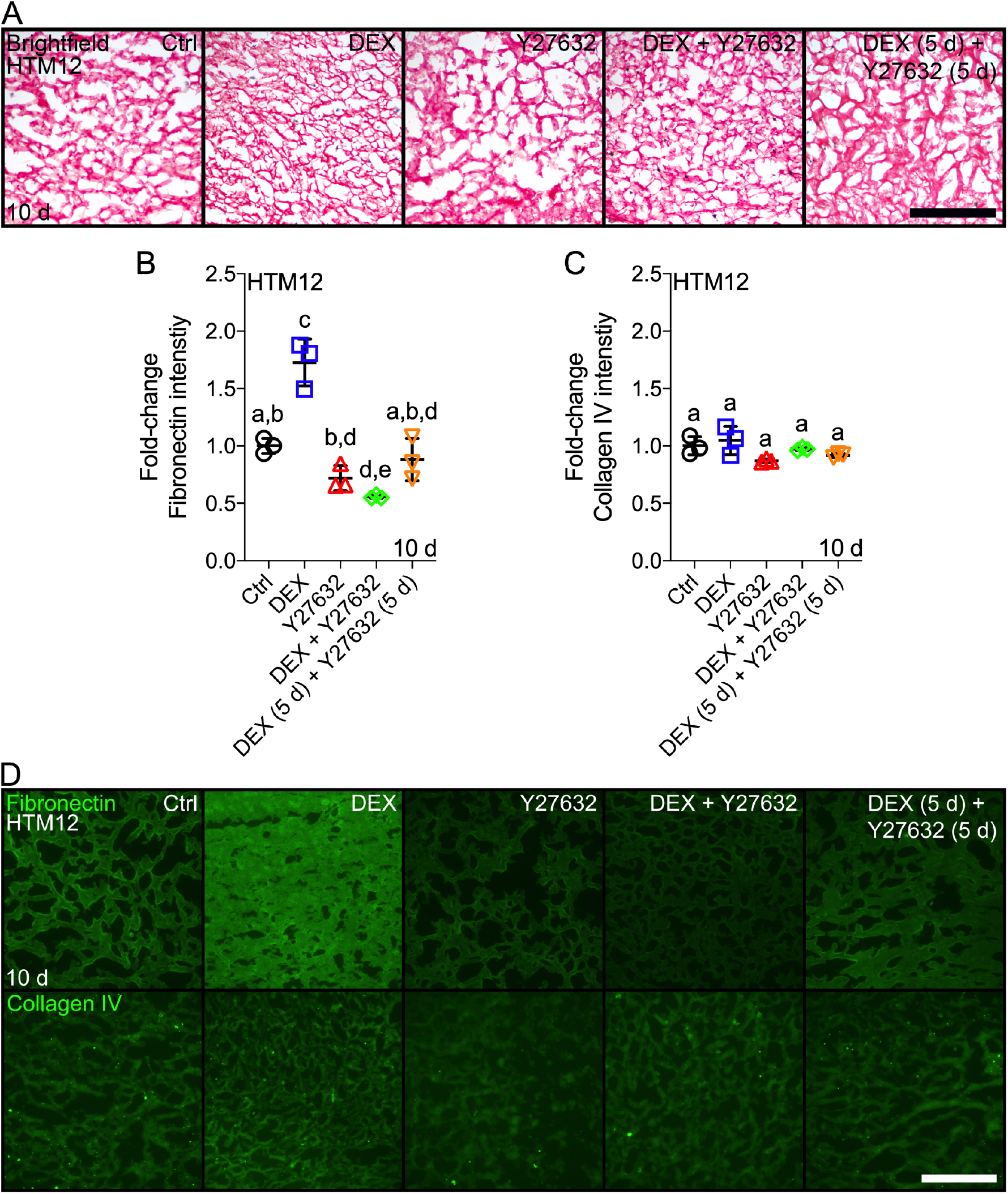
HTM hydrogel morphology and ECM deposition with corticosteroid induction and ROCK inhibitor rescue. (**A**) Representative brightfield micrographs of picrosirius red/hematoxylin-stained sections from HTM12 hydrogels subjected to control, 100 nM DEX, 10 µM Y27632, DEX + Y27632, or DEX (5 d) + Y27632 (5 d) at 10 d (hydrogel/ECM = pink; HTM cells = blue). Scale bar, 250 μm (**B,C**) Quantification and (**D, E**) representative fluorescence micrographs of fibronectin and collagen IV (= green) in HTM12 hydrogels subjected to the different treatments at 10 d. Scale bar, 250 μm.

There is increased accumulation of ECM proteins in the HTM tissue of glaucomatous eyes ^66^. Therefore, we next evaluated HTM cell-mediated fibronectin and collagen type IV deposition within hydrogels in response to the different treatments (**Fig. 9B-D**). DEX-treated constructs exhibited significantly more fibronectin deposition vs. controls (highest overall), whereas Y27632 markedly reduced the fibronectin signal intensity. DEX + Y27632 co-treatment resulted in significantly less fibronectin deposition (lowest overall) compared to the control and DEX groups. Importantly, Y27632 rescued aberrant fibronectin deposition induced by DEX; i.e., significantly less signal intensity with DEX (5 d) + Y27632 (5 d) sequential treatment was noted compared to DEX-treated samples (**Fig. 9B,D**). Similar trends (albeit not significant) were observed for collagen IV, with overall lower expression compared to fibronectin (**Fig. 9C,D**).

Together, these data show that DEX induces HTM hydrogel condensation and remodeling, and aberrant ECM deposition consistent with glaucomatous tissue dysfunction, which is largely rescued with ROCK inhibition.

### Hydrogel stiffness

Dysregulation of HTM cells and pathologic ECM remodeling jointly contribute to HTM stiffening in POAG, which negatively affects IOP in a feed-forward loop ^11–13^. To assess the functional consequences of increased hydrogel contraction/condensation, driven by cytoskeletal rearrangements and ECM remodeling/deposition, on tissue-level construct stiffness, we next performed rheology analysis of HTM hydrogels subjected to the different treatments. We determined fold-changes in storage modulus vs. 0 d for comparisons between groups (**Fig. 10A**). DEX treatment significantly stiffened HTM hydrogels vs. control (~1.7-fold) consistent with recent reports from normal and glaucomatous HTM tissues ^13,67^, while Y27632 had precisely opposite effects and significantly softened HTM hydrogels vs. controls (~0.6-fold). No differences were observed between DEX + Y27632 co-treatment and controls. Importantly, Y27632 rescued pathologic HTM hydrogel stiffening induced by DEX and prevented further stiffening; i.e., storage modulus with DEX (5 d) + Y27632 (5 d) sequential treatment was similar to the control and DEX + Y27632 groups, but significantly lower compared to DEX-treated samples (**Fig. 10A**). These findings were in agreement with HTM hydrogel contraction in this large sample format (**Fig. 10B,C**), mirroring the trends observed with the 15x smaller samples (**Fig. 6C**).

**Fig. 10.**
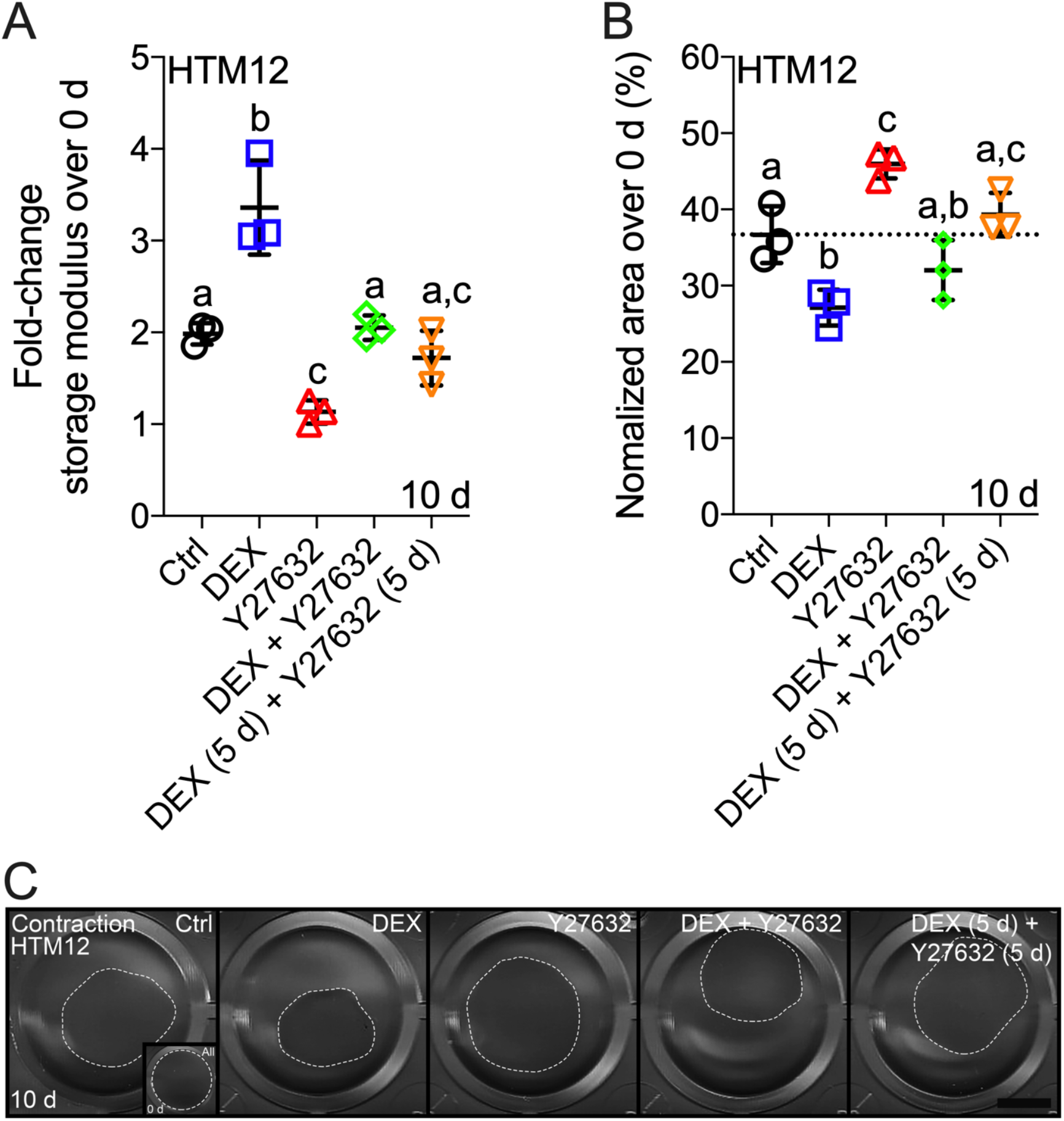
HTM hydrogel stiffness with corticosteroid induction and ROCK inhibitor rescue. (**A**) Fold-change in storage modulus (i.e., normalized to 0 d) of HTM12 hydrogels subjected to control, 100 nM DEX, 10 µM Y27632, DEX + Y27632, or DEX (5 d) + Y27632 (5 d) at 10 d (N=3 per group; shared significance indicator letters represent non-significant difference (p>0.05), distinct letters represent significant difference (p<0.05)). (**B**) Normalized contractility quantification (i.e., construct size relative to 0 d) of HTM12 hydrogels subjected to the different treatments at 10 d (N=3 per group; dotted line shows control values for reference; shared significance indicator letters represent non-significant difference (p>0.05), distinct letters represent significant difference (p<0.05)). (**C**) Representative scanned 24-well plate with HTM12 hydrogels subjected to the different treatments at 10 d (white dashed lines outline size of contracted constructs; inset, white dashed line outlines original size of all constructs [= entire well area] at 0 d). Scale bar, 5 mm.

This suggests that DEX induces tissue-level HTM hydrogel contraction and stiffening consistent with glaucomatous tissue behavior, and that ROCK inhibition effectively prevents or rescues DEX-induced pathologic stiffening.

## Discussion

*In vitro* studies of HTM cell function are fundamentally limited by traditional cell culture systems. It is widely accepted that cells in 3D environments made of relevant ECM proteins show altered behavior compared to 2D, highlighting the need for more biologically-relevant HTM model systems ^68,69^. Current bioengineered models ^37, 39–41, 47–49,51,70^ are useful tools for investigating aspects of HTM cell/tissue behavior under normal and simulated glaucomatous conditions, and complement proven *ex vivo* perfusion-cultured anterior segments and *in vivo* animal models. However, they cannot accurately mimic the complex native HTM cell-ECM interface, which makes them less than ideal to investigate the processes involved in the onset and progression of glaucomatous HTM stiffening typically observed with POAG. In this study, we used ECM proteins present in the native tissue to formulate a novel biomimetic 3D HTM hydrogel to provide a key advancement in bioengineered glaucoma disease modeling.

Our focus was to partially recapitulate the JCT region for both its critical role in regulating AH outflow resistance and IOP, and inextricable link to glaucomatous HTM tissue stiffening. HTM cells within the JCT are surrounded by fibrillar ECM components (i.e., collagen type I, elastin) to form a loose connective tissue rich in the glycosaminoglycan HA ^3–7^. Proper maintenance of this HTM cell-ECM microenvironment is critical for homeostatic IOP regulation via normal ECM remodeling ^4–7^. Protein-based hydrogels are widely used in tissue engineering applications as they provide a simplified version of the natural 3D tissue environment and allow for accurate *in vitro* modeling of cellular behaviors ^42–46^. Photoactive functional groups (e.g., thiols, methacrylates) can be chemically conjugated to protein-based materials. These functionalized precursors can then be crosslinked by exposure to UV light in the presence of a biocompatible photoinitiator, without or with cells, to form stable hydrogel networks with high water content ^46^. Collagen type I and HA are frequently used for tissue engineering applications and functionalized polymers are commercially available; however, incorporating elastin presents certain challenges. In native tissues, elastic fibers are highly crosslinked and insoluble preventing their direct processing for use in biomaterials ^71^. Therefore, various forms of soluble elastin have been engineered, including ELPs ^72,73^. The key advantage of the unique ELP used herein is that it is flanked by a short peptide sequence (K<u>C</u>TS) in which the two cysteines effectively act as photoactive thiols ^56^. A UV-crosslinked hydrogel containing ELP and HA was recently reported ^57^; however, no study to date has developed a biopolymer composite system with added collagen type I, the most abundant fibrillar protein in the human body and main structural element of the ECM ^74^.

We formulated a novel HTM hydrogel by mixing normal human donor-derived TM cells with methacrylate-conjugated collagen type I, KCTS-flanked ELP, and thiol-conjugated HA followed by photoinitiator-mediated short UV crosslinking. Informed by previous studies ^75^, we showed excellent HTM cell viability (i.e., negligible cytotoxicity) in response to 5 s UV crosslinking at the chosen conditions. This protein-based hydrogel created a supportive microenvironment to facilitate prominent HTM cell spreading/proliferation throughout the adequately soft and porous 3D network. Of note, we estimated (acellular) hydrogel pore size at ~40 µm, which is larger than that found in the JCT region but consistent with the central and outer lamellar regions ^4,5,7^. Higher concentrations of biopolymer precursors could provide increased crosslinking density, which may increase the hydrogel stiffness and consequently decrease pore size. Dynamic interactions between HTM cells and their surrounding ECM resulted in hydrogel condensation/contraction and remodeling, mimicking critical normal tissue behavior. As such, our HTM cell-encapsulated hydrogel containing (by weight) ~68% collagen type I, ~26% ELP, and ~5% HA is the first of its kind with distinct biomimetic advantages over other HTM model systems ^37, 39–41, 47–49,51,70^.

Prolonged treatment with ocular corticosteroids such as DEX can lead to steroid-induced elevation of IOP. This in turn can cause steroid-induced glaucoma, which has commonalities with POAG; namely, increased HTM cell/tissue contraction and ECM deposition that can lead to HTM stiffening and increased resistance to AH outflow ^9,10,76,77^. Therefore, DEX is frequently used to induce glaucomatous conditions in experimental studies ^35^. Rho-associated kinases are effectors of the small GTPase Rho, which play major roles in mediating the reorganization of the actin cytoskeleton ^78,79^ with involvement in diseases such as pulmonary hypertension, nerve injury, and POAG ^80^. To that end, the recently approved ROCK inhibitor netarsudil ^21^ directly targets the stiffened HTM to increase AH outflow via cell/tissue relaxation through reduction of actomyosin contractile tone ^22–26^. And while not without merit, current HTM model systems are hampered by their inability to reliably and efficiently determine the relative contributions of HTM cells and their ECM to the onset and progression of glaucomatous HTM stiffening. No *in vitro* study to date has reported changes in tissue-level construct stiffness with DEX challenge/Y27632 rescue. To assess the utility of our HTM hydrogel as glaucoma disease model, we used proven corticosteroid induction of POAG-like conditions with DEX, and assessed the therapeutic effects of ROCK inhibition with Y27632 on cell cytoskeletal organization and tissue-level function dependent on HTM cell-ECM interactions.

We showed increased HTM hydrogel contraction with DEX treatment, which is opposite to what a previous study found using simple collagen type I hydrogels encapsulated with bovine TM cells ^81^, but consistent with data obtained from perfusion-cultured HTM tissues challenged with DEX ^76,77^. Actin abundance and organization are thought to modulate intracellular force generation and cell contractility; our results showed that DEX induced elevated f-actin and αSMA abundance and rearrangement, which is consistent with observations made using porous scaffolds seeded with HTM cells ^38^. This suggests that increased actin abundance could induce cell contractile force generation to condense HTM cells within the 3D hydrogel network resulting in overall increased construct contractility. HTM remodeling is critical for proper tissue function, driven by the reciprocity between HTM cells and their ECM. By the same token, pathologic ECM remodeling contributes to glaucomatous tissue dysfunction. We observed increased microstructural ECM remodeling (i.e., condensed hydrogels with decreased inter-fibrillar spaces and reduced fiber thickness) with DEX treatment. Furthermore, fibronectin - and to a lesser extent collagen type IV - deposition induced by DEX was confirmed in our model, which has been reported to contribute to elevated outflow resistance in the HTM with POAG ^9^. These findings are in agreement with glaucomatous pathology and have all been linked to progressive HTM tissue stiffening. Importantly, consistent with the reported reduction of actomyosin contraction in HTM tissue following ROCK inhibition ^22–25^, we showed that Y27632 co-treatment (simulating a prophylactic approach) and sequential treatment (simulating a therapeutic approach) potently rescued the DEX-induced pathologic changes in HTM hydrogel contractility, condensation, and ECM deposition/remodeling. Using an alternative pair of pharmacological induction and rescue of glaucomatous conditions - TGF-β2 (a potent cytokine implicated in POAG pathology ^63^) and Lat-B (an actin filament depolymerizing agent ^64^), we were able to validate the utility of our HTM hydrogel as glaucoma disease model.

Patients with POAG, the most common subtype of glaucoma, typically suffer from elevated IOP and experience increased HTM stiffness, which in turn can negatively affect IOP and HTM cell function in a feed-forward loop ^9–13^. It is perhaps because of these observations that more studies in recent years have focused on the biomechanical properties of tissues within the conventional outflow pathway, and their dynamic changes in disease. HTM tissue stiffness has been determined directly and indirectly in several studies. In an early report, HTM stiffness was shown to be ~20-fold higher in glaucomatous vs. normal eyes by atomic force microscopy (AFM) ^11^. Several limitations to this study, however, may have resulted in overestimated differences in tissue elasticity. In a recent elegant study using a novel indirect approach (with AFM correlation), HTM stiffness was determined to be ~1.4-fold greater in glaucomatous HTM tissues vs. normal controls ^13,67^. The discrepancies in both absolute and relative stiffness values (i.e., normal vs. glaucomatous) between the two studies most likely stem from methodological differences. We here showed an ~1.7-fold increase in HTM hydrogel stiffness with DEX treatment vs. controls as measured by rheology (oscillatory shear-strain sweep test), a common technique for characterizing biomechanical properties of viscoelastic hydrogels. Y27632 treatment had precisely opposite effects, and was shown to prevent or rescue the DEX-induced pathologic changes, resulting in HTM hydrogel softening. This was in agreement with the HTM hydrogel contraction, cytoskeletal rearrangement, and ECM remodeling/deposition data suggesting potent therapeutic potential of Y27632 in support of clinical netarsudil use. While we can only speculate on the corresponding hydrogel stiffness values by AFM, previous studies have suggested good overall correlation between the two methods ^82,83^, with trends frequently being preserved.

Elevated AH outflow resistance and IOP are directly related to retinal ganglion cell damage in POAG. We acknowledge that a limitation to our system is the lack of flow. Ongoing work to integrate the HTM hydrogel, with addition of a Schlemm’s canal endothelial cell layer, on a microfluidics platform for measurements of flow resistance under dynamic conditions is expected to further expand the utility of our hydrogel-based glaucoma disease model. ECM stiffness is considered a universal mechanical cue that can both precede and propel disease by altering the behavior of associated cells according to the changing ECM elasticity. In diseases such as idiopathic lung fibrosis and multiple myeloma, there are clinical trials underway that directly target ECM remodeling to prevent pathologic tissue stiffening ^30^. If matrix stiffness is a driving factor that causes the ECM to further stiffen under pathological conditions, then what are the initial mechanisms contributing to pre-pathological increases in matrix rigidity? A key advantage of our HTM hydrogel is the ability to tune construct stiffness. Ongoing work explores the potential of targeting relevant mechanotransduction pathways for POAG therapy.

In sum, we have developed a novel *in vitro* biomimetic HTM hydrogel model that allows for detailed investigation of the 3D HTM cell-ECM interface under normal and simulated POAG conditions upon *in situ* corticosteroid induction. Importantly, unlike other model systems, the HTM biopolymer hydrogel enables correlative analyses of HTM cell cytoskeletal organization with tissue-level functional changes - glaucomatous pathologic contraction and stiffening - contingent on HTM cell-ECM interactions. The bidirectional responsiveness of the HTM hydrogel model system to pharmaceutical challenge and rescue suggests promising potential to serve as a screening platform for new POAG treatments with focus on HTM cell/tissue biomechanics.

## Acknowledgments

We thank Iris Navarro at the Duke Eye Center - BioSight Program for guidance in setting up initial HTM isolation and characterization procedures. We also thank the team at Specialty Surgery Center of Central New York for assistance with corneal rim specimens. We thank Dr. Audrey M. Bernstein, Dr. Mariano S. Viapiano, and Dr. Jason A. Horton for imaging support. We also thank the teams at the Syracuse University - Syracuse Biomaterials Institute and BioInspired Institute for access to shared facilities and technical advice. We thank Benjamin Zink and Debra Driscoll at the SUNY College of Environmental Science and Forestry - Analytical and Technical Services for assistance with SEM imaging.

## Author contributions

H.L., A.E.P., P.S.G., and S.H designed all experiments, collected, analyzed, and interpreted the data. T.B. and A.K. assisted with HTM cell isolation, ELP expression/purification, PDMS-mold design/3D printing, and imaging. R.W.W., N.A., and W.D.S. provided study materials. H.L. and S.H. wrote the manuscript. All authors collected data and commented on and approved the final manuscript. P.S.G. and S.H. conceived and supervised the research.

## Data and materials availability

All data needed to evaluate the conclusions in the paper are present in the paper and/or the Supplementary Materials. Additional data related to this paper may be requested from the authors.

