## Supplemental information for "A tissue-engineered human trabecular meshwork hydrogel for advanced glaucoma disease modeling"

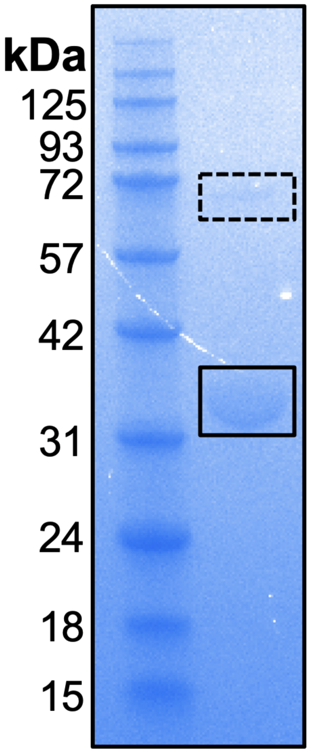


**Suppl. Fig. S1. ELP characterization.** Coomassie blue-stained protein electrophoresis gel of ELP showed 32.5 kDa (solid box), and a weak band at approximately double the molecular weight (dashed box).

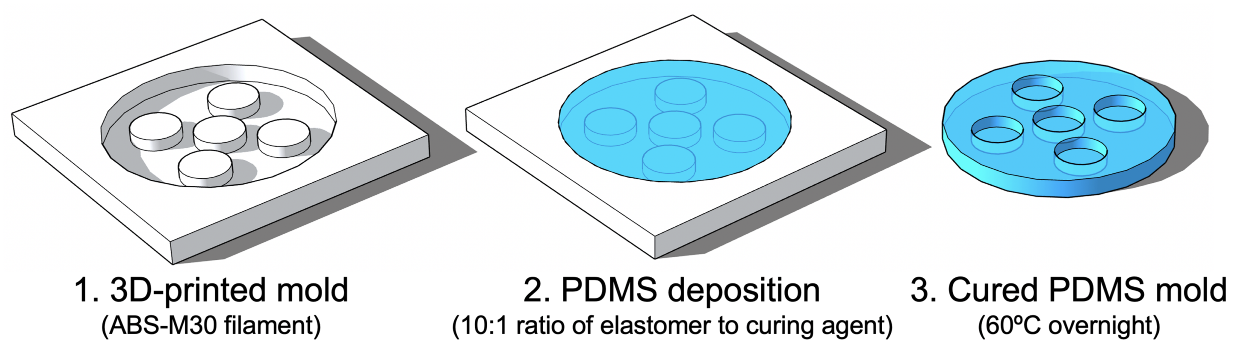


**Suppl. Fig. S2. PDMS mold fabrication.** Custom master molds (8x1-mm, 8x2-mm) were 3D-printed using ABS-M30 filament. Polydimethylsiloxane (PDMS) was mixed as per the manufacturer’s protocol, poured into the 3D-printed negative master molds, degassed under vacuum, and cured overnight at 60ºC.

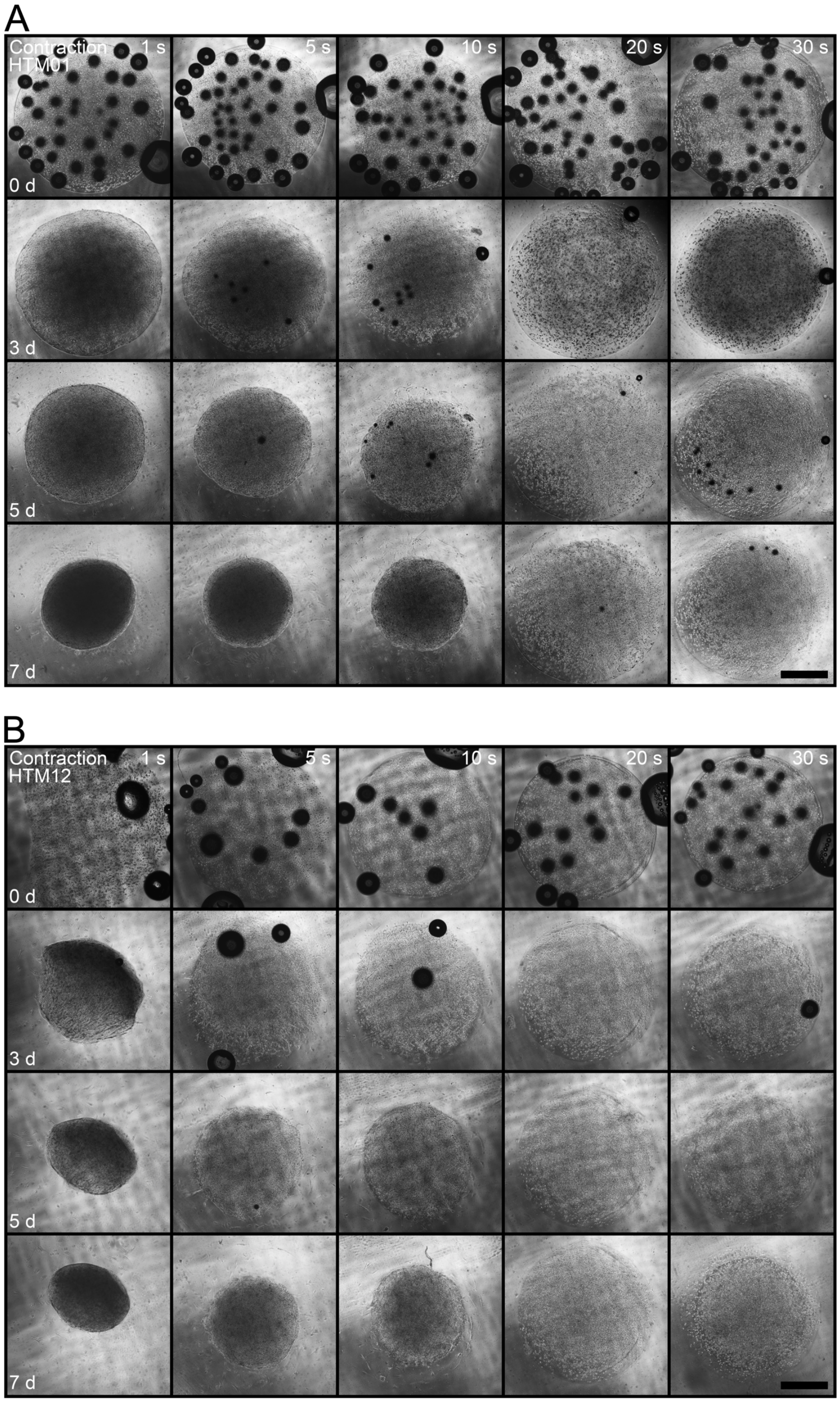


**Suppl. Fig. S3. HTM hydrogel contractility.** Representative brightfield images of (**A**) HTM01 and (**B**) HTM12 hydrogels across UV crosslinking times at 0, 3, 5 and 7 d. Scale bars, 1 mm.

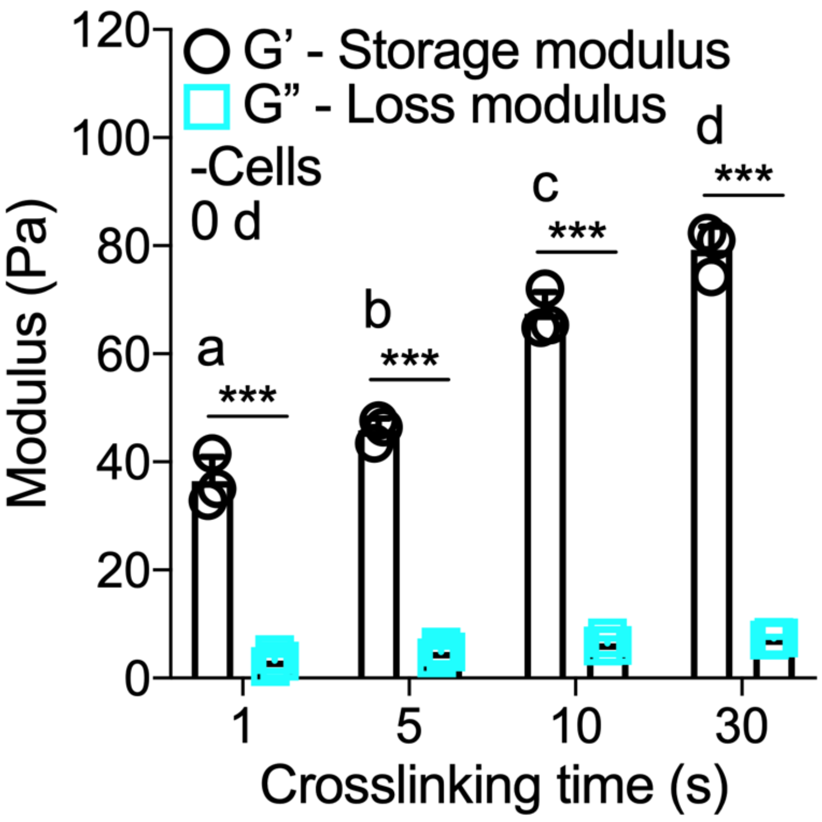


**Suppl. Fig. S4. Hydrogel mechanical analysis.** Storage (G') and loss moduli (G") of acellular hydrogels UV-crosslinked for 1, 5, 10 or 30 s at 0 d (N=3 per group; shared significance indicator letters represent non-significant difference ($p>0.05$), distinct letters represent significant difference ($p<0.05$)).

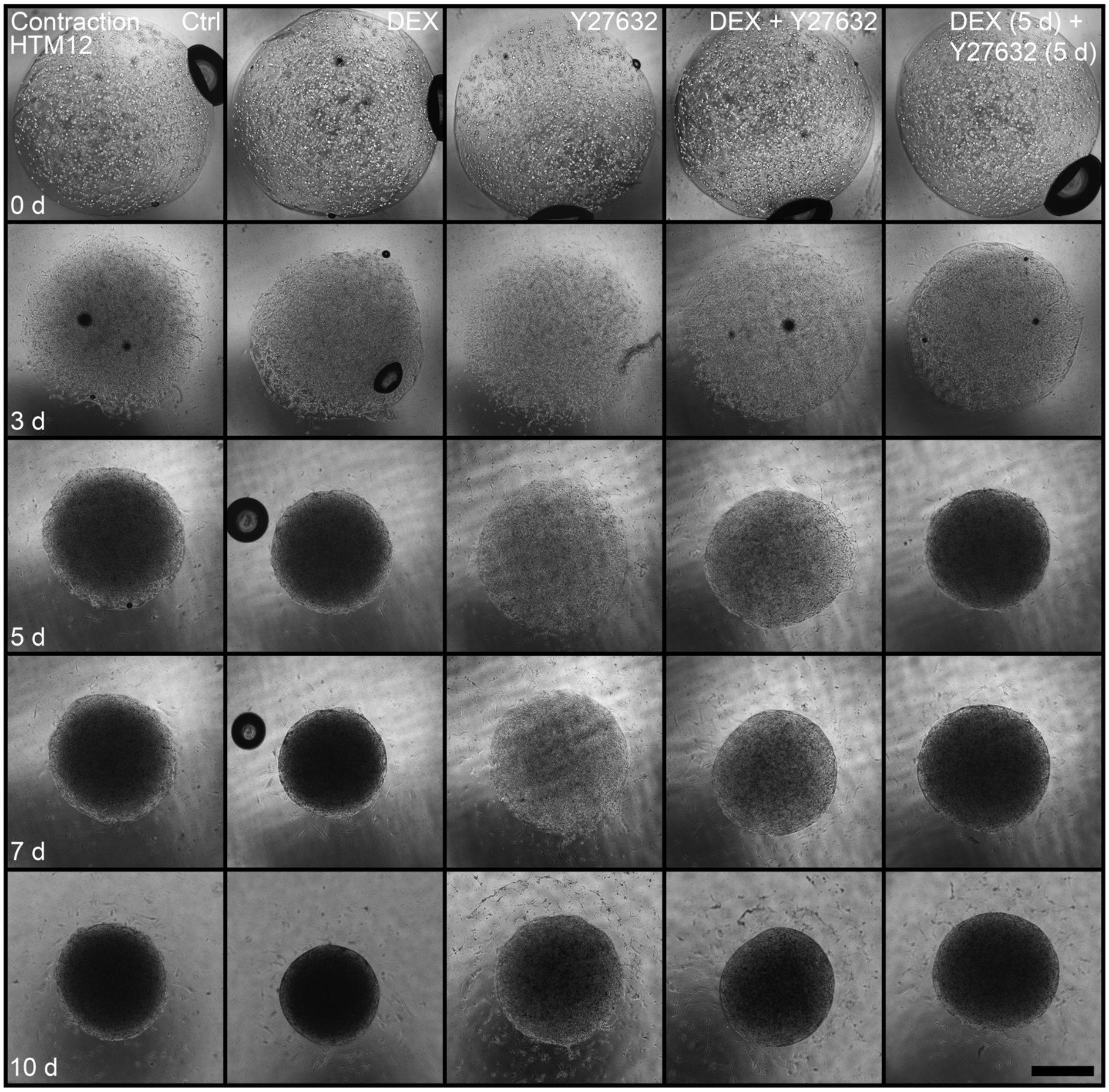


**Suppl. Fig. S5. HTM hydrogel contractility with corticosteroid induction and ROCK inhibitor rescue.** Representative brightfield images of HTM12 hydrogels subjected to control, 100 nM DEX, 10 μM Y27632, DEX + Y27632, or DEX (5 d) + Y27632 (5 d) at 0, 3, 5, 7 and 10 d. Scale bars, 1 mm.

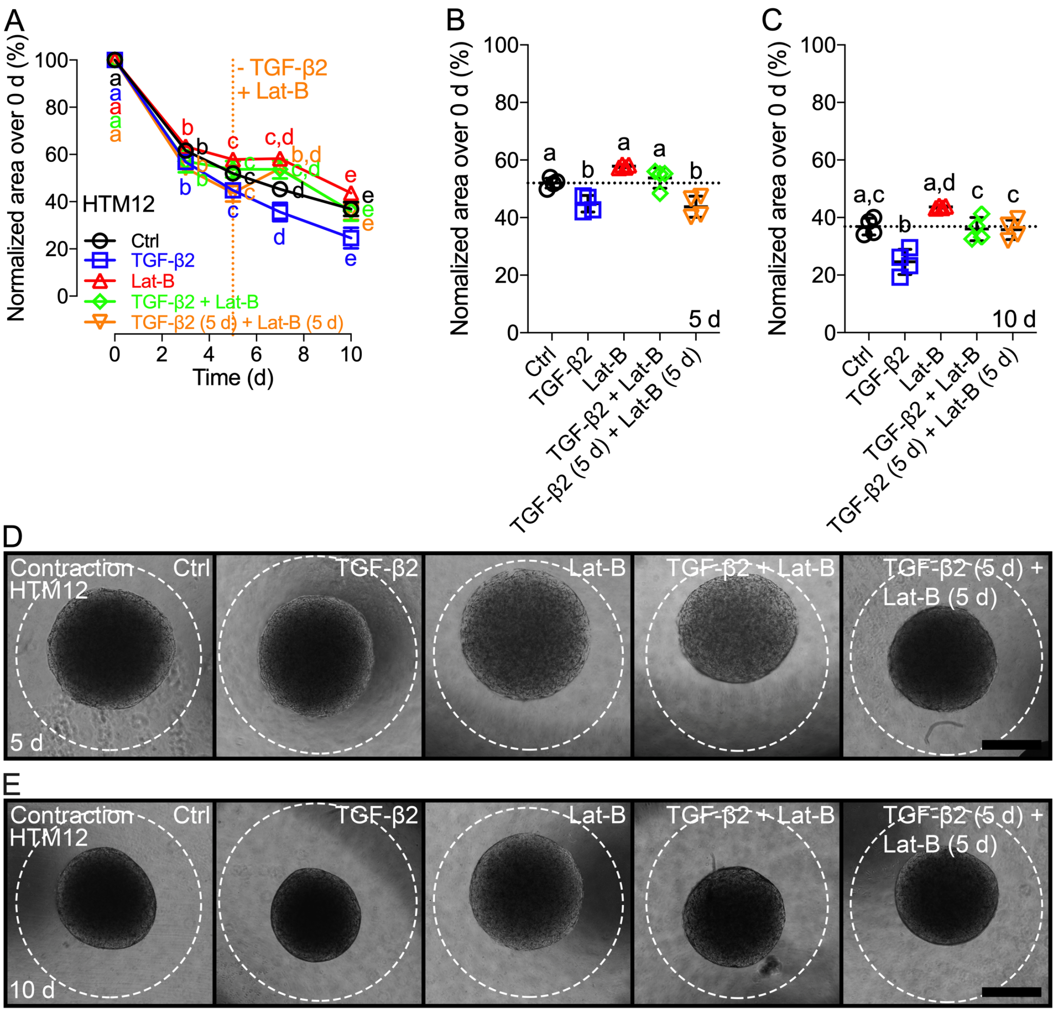


**Suppl. Fig. S6. HTM hydrogel contractility with cytokine induction and actin depolarization rescue.** (**A**) Longitudinal quantification (i.e., construct size relative to 0 d) of HTM12 cell contractility in hydrogels subjected to control, 2.5 ng/ml transforming growth factor beta-2 (TGF-β2), 2 μM latrunculin-b (Lat-B), TGF-β2 + Lat-B, or TGF-β2 (5 d) + Lat-B (5 d) (N=4 per group; shared significance indicator letters represent non-significant difference ($p>0.05$), distinct letters represent significant difference ($p<0.05$)). Detailed comparisons between groups at (**B**) 5 d and (**C**) 10 d (dotted lines show respective control values for reference). Representative brightfield images of HTM12 hydrogels subjected to the different treatments at (**D**) 5 d and (**E**) 10 d; white dashed lines outline original size of constructs at 0 d. Scale bars, 1 mm.
